# NKTR-255, a polymer-conjugated IL-15, synergizes with CAR-T cell therapy to activate endogenous anti-tumor immunity and improve tumor control

**DOI:** 10.64898/2025.12.11.693768

**Authors:** W. Sam Nutt, Mitchell G. Kluesner, Emma Bingham, Ekram Gad, Dani Miller, Victor Zepeda, Andrew J. Snyder, Sydney A. Marsh, Shannon M. Liudahl, Mari Hoffman, Lillian LeBlanc, Madeleine E. Volfbeyn, Sarah M. Garrison, Megha Sarvothama, Kevin C. Barry, Mark B. Headley, Mario Marcondes, Shivani Srivastava

**Affiliations:** Human Biology Division, Fred Hutchinson Cancer Center; Seattle, WA, USA; Molecular and Cell Biology Graduate Program, University of Washington; Seattle, WA, USA; Medical Scientist Training Program, University of Washington School of Medicine; Seattle, WA, USA; Comparative Medicine (TRMS), Fred Hutchinson Cancer Center; Seattle, WA, USA; Translational Science and Therapeutics Division, Fred Hutchinson Cancer Center; Seattle, WA, USA; Immunology Graduate Program, University of Washington; Seattle, WA, USA; Public Health Sciences Division, Fred Hutchinson Cancer Center; Seattle, WA, USA; Nektar Therapeutics; San Francisco, CA, USA

## Abstract

CAR-T cells have yet to show widespread efficacy in solid tumors due in part to their poor persistence and loss of function in the tumor microenvironment. Further, heterogenous expression of most CAR target antigens in solid tumors can lead to escape of antigen-null tumors that resist CAR-T killing. Strategies to cooperatively boost both CAR-T and endogenous anti-tumor immunity could curb tumor escape and may be critical for achieving durable efficacy in cancer patients. NKTR-255 is a polymer-conjugated IL-15 with extended half-life that can boost endogenous T and NK cells, as well as CD19 CAR-T activity in B cell malignancies. However, whether NKTR-255 is sufficient to overcome CAR-T dysfunction in the suppressive solid tumor microenvironment, and how NKTR-255 and CAR-Ts together re-shape endogenous anti-tumor immunity, is not known. Using an autochthonous mouse model of ROR1^+^ lung adenocarcinoma, we show that NKTR-255 significantly boosted accumulation, reduced exhaustion, and improved function of tumor-infiltrating CAR-T cells. Compared with NKTR-255 or CAR-T treatment alone, combination of NKTR-255 and CAR-T therapy synergistically increased tumor-infiltrating CD11b^+^ cytotoxic NK cells, activated dendritic cells, and endogenous tumor-specific T cells that preserved a PD-1^+^Tcf1^+^ stem-like phenotype. Consequently, NKTR-255 and CAR-T combination therapy induced complete elimination of ROR1^+^ tumor and significantly improved survival, with enhanced tumor control dependent on activity of both CAR-Ts and endogenous T cells. Altogether, our data suggest that combining NKTR-255 with CAR-T therapy is a promising strategy to enhance both CAR-T and endogenous anti-tumor immunity to promote coordinated control of aggressive tumors.

## Introduction

CAR-T cell therapy has mediated remarkable responses in patients with hematologic malignancies, resulting in FDA approval of multiple CAR-T cell products targeting CD19 or BCMA.^1^ While recent studies have described clinical responses to CAR-T therapy in more common solid tumors, response rates are low, and CAR-T therapy remains widely ineffective for most patients with solid tumors.^2^ Several studies suggest that CAR-T efficacy in solid tumors is limited by poor persistence and dysfunction of infused cells, as well as heterogeneous expression of most CAR target antigens.^3^ Indeed, in a phase I trial conducted at our Center, CAR-T cells targeting the tumor-associated antigen ROR1 induced complete responses in patients with chronic lymphocytic leukemia (**CLL**) but rapidly lost function and persisted poorly in patients with triple-negative breast cancer (**TNBC**) and non-small cell lung cancer (**NSCLC**).^4,5^ Moreover, in patients who transiently respond to CAR-T therapy, efficacy is limited by progression of tumor cells that lack or downregulate expression of the CAR target.^4,6–15^

One strategy to overcome tumor escape is to engage endogenous immune cells to target tumor cells that have downregulated the CAR target, leading to a broader and more effective anti-tumor response.^16,17^ One axis of endogenous immune activation is “epitope spreading,” which occurs when a T cell specific for one tumor antigen induces tumor cell death, resulting in uptake and presentation of additional tumor antigens by dendritic cells (**DCs**), and subsequent activation of T cells specific for additional tumor antigens.^16,17^ CAR-T cells may be well-suited to initiating this process due to their robust ability to induce tumor cell death, antigen release, and secretion of pro-inflammatory cytokines that can activate both DCs and endogenous T cells.^18,19^ In fact, epitope spreading has been observed clinically following adoptive T cell therapy in some patients and is associated with improved outcomes.^20–23^ Thus, strategies to cooperatively boost both CAR-T and endogenous anti-tumor immunity may be critical for achieving durable efficacy in patients.

IL-15 is a critical cytokine that promotes T cell proliferation, survival, and effector function.^24–28^ Consistent with this, increased serum IL-15 levels prior to CAR-T cell infusion correlate with greater peak CAR-T cell expansion in patients with hematologic or solid tumors, suggesting that administering exogenous IL-15 could further boost CAR-T cell activity.^4,29^ Compared to the related cytokine IL-2, which signals through the same IL-2Rβ/IL-2Rγ receptor complex, IL-15 has been shown to preferentially promote proliferation and self-renewal of a stem-like subset of PD-1^+^Tcf1^+^ precursor exhausted cells (**Tpex**),^30^ as well as promote survival of the effector-like cells they generate,^31^ suggesting it may be well-suited to overcome CAR-T and endogenous T cell dysfunction in the solid tumor microenvironment.^26,32^ While multiple drugs targeting the IL-15/IL-15R axis are in active development,^26,28,32^ one of the leading candidates advanced to clinical testing is NKTR-255, a polymer-conjugated IL-15 that shows extended half-life and boosts T cells and NK cells more effectively than recombinant IL-15.^33,34^ NKTR-255 has shown promising results in combination with CD19-targeting CAR-T cells in patients with B cell malignancies,^29,35–37^ highlighting its safety and efficacy in enhancing peak CAR-T cell expansion. However, it is not known whether NKTR-255 is sufficient to overcome CAR-T dysfunction and exhaustion within the suppressive microenvironment of solid tumors like lung cancer. Additionally, as most studies of NKTR-255 and CAR-T therapy have been performed in xenograft tumor models in immunodeficient mice,^29^ it is unknown how NKTR-255 and CAR-T cell therapy together affect endogenous anti-tumor responses, how these changes contribute functionally to tumor control, and whether this combination strategy provides distinct benefits beyond either monotherapy.

We previously developed an autochthonous model of ROR1^+^ lung adenocarcinoma that mimics the initiation, progression, and suppressive tumor microenvironment of human NSCLC, as well as the poor persistence and dysfunction of ROR1 CAR-T cells observed in NSCLC patients, providing a unique platform to test strategies to enhance both CAR-T and endogenous anti-tumor immunity that can inform clinical translation.^5,38,39^ Here, we use transplantable and autochthonous models of ROR1^+^ lung cancer to investigate how NKTR-255 affects ROR1 CAR-T cell activity and evaluate how NKTR-255 and CAR-T treatment together reshape endogenous anti-tumor immunity.

## Materials & Methods

### EXPERIMENTAL MODEL AND SUBJECT DETAILS

#### Animals

C57BL/6 (**B6**), B6.SJL (**B6 CD45.1**), and B6 Thy1.1 mice were purchased from Jackson Laboratory. *Kras^LSL-G12D/+^; p53^fl/fl^* (**KP**) mice were generously provided by A. McGarry Houghton (Fred Hutch Cancer Center (**FHCC**), Seattle WA). For studies with KP mice, 8-18 week old age-matched and sex-matched mice were used. For all other studies, 8-12-week-old age-matched and sex-matched mice were used. Mice of the same sex were randomly assigned to experimental groups or were assigned based on tumor burden, such that average tumor burden was consistent across experimental groups prior to treatment. All mice were housed and bred at the FHCC (Seattle, WA). All experiments were performed in accordance with the guidelines of the FHCC Institutional Animal Care and Use Committee (PROTO202000031).

#### Cell Lines

The KP1233 (“**KP**”) tumor and 3TZ “GreenGo” cell lines were generously provided by Tyler Jacks (Massachusetts Institute of Technology). Lenti-X cells for lentiviral packaging were purchased from Clontech/Takara (#632180). Lenti-X and 3TZ were maintained in complete DMEM (cDMEM = DMEM supplemented with 10% FBS, 2 mM L-glutamine, 100 U/ml penicillin/streptomycin, 25 mM HEPES). Platinum-E (Plat-E) cells for retroviral packaging were purchased from Cell Biolabs and were maintained in cDMEM with added 1ug/mL puromycin and 10ug/mL blasticidin. The KP^ROR1^ tumor cell line was generated by retroviral transduction of the KP tumor cell line with full-length human *ROR1* cDNA (UniProt: <u>Q01973</u>) and subsequent FACS sorting of ROR1^+^ cells to >95% purity, as previously described.^5^ To generate the KP^ROR1-Ova^ cell line, the retroviral vector was modified to encode truncated human ROR1 (UniProt: Q01973, aa1-462) linked by a P2A skip element to truncated CD19 (UniProt: P25918, aa1-321) fused via a SSSSG linker to the MHC class I-restricted epitope, SIINFEKL, derived from chicken ovalbumin (UniProt: P01012, aa249-271). KP tumor cells were retrovirally transduced to express the ROR1-Ova construct, and ROR1^+^CD19^+^ tumor cells were FACS sorted to >95% purity. KP^ROR1^ and KP^ROR1-Ova^ were maintained in cDMEM. All cells were tested bi-monthly for the absence of mycoplasma.

### METHOD DETAILS

#### Cloning of Murine CAR Constructs

The MP71 retroviral vector was modified to encode a human ROR1-specific CAR co-expressed with a truncated murine CD19 transduction marker (**tCD19**), as described.^5^ In some experiments, the tCD19 transduction marker was fused to eGFP (tCD19/GFP). The ROR1 CAR possessed a murine CD8α signal peptide (UniProt: P01731, aa1-27), 2A2 scFv,^5,40^ human IgG4 short spacer,^41^ murine CD28 transmembrane (UniProt: P31041, aa151-177), murine 4-1BB signaling domain (UniProt: P20334, aa211-256), murine CD3ς (UniProt: P24161, aa52-164), and was linked by a P2A ribosomal skip element to tCD19 or tCD19/GFP.

#### Generation of Murine CAR-T Cells

Retrovirus was produced by transient transfection of Plat-E cells with the indicated MP71 vectors using the CalPhos Mammalian Transfection Kit (Takara). Viral supernatant was harvested 48 hr after transfection, filtered through a 0.45 μm pore filter, and snap-frozen in liquid nitrogen for long-term storage. T cell suspensions were prepared from spleen and peripheral lymph nodes of donor mice by using 1mL syringe plungers to mechanically dissociate tissues through a 40 μm filter. Murine CD8^+^ and CD4^+^ T cells were enriched from spleens and peripheral lymph nodes of donor mice using untouched negative isolation kits (EasySep Mouse CD4^+^ T Cell Isolation Kit: #19852, EasySep™ Mouse CD8^+^ T Cell Isolation Kit: #19853) and stimulated with anti-CD3/28 Mouse T-Activator Dynabeads (ThermoFisher: #11453D) at a bead to cell ratio of 1:1 for 24 hr in a 37°C, 5% CO_2­­_incubator in complete RPMI (RPM1 1640, 10% heat inactivated FBS, 1 mM sodium pyruvate, 1 mM HEPES, 100 U/ml penicillin/streptomycin, 50 μM β-mercaptoethanol) supplemented with 50 U/ml recombinant human IL-2 (Peprotech #200-02). Non-tissue culture plates were coated with 12.5 μg/ml RetroNectin (Takara: #T100B) according to the manufacturer’s protocol, and plates were loaded with pre-titered retrovirus and centrifuged for 2 hr at 3000xg at 32°C. Murine T cells after 24 hr incubation were harvested and resuspended at 1×10^6^ cells/ml in fresh complete RPMI supplemented with 50 U/ml IL-2. Viral supernatant was aspirated from RetroNectin-coated plates, plates were rinsed with PBS, and T cells were added to each virus-coated well. Plates were then centrifuged at 800g for 30 min at 32°C and returned to 37°C, 5% CO_2­_incubators. After 24 hours, T cells were harvested, counted and resuspended in complete RPMI with 50 U/ml IL-2 to a concentration of 0.5x10^6^ cells/mL. T cells were subsequently harvested, counted, and maintained at 0.5×10^6^ cells/ml in complete RPMI with 50 ng/ml human IL-15 for two days after. Four days after transduction, magnetic beads were removed and T cell transduction was measured by flow cytometry staining for tCD19, GFP, and ROR1 CAR. Transduced cells were enriched using the EasySep CD19 Positive Selection Kit II (Stem Cell: #18954) and either used immediately or cryopreserved in complete RPMI supplemented with 10% FBS and 10% DMSO. Cryopreserved cells were thawed in a 37°C water bath, washed, counted, re-confirmed for viability, tCD19, GFP, and ROR1 CAR expression by flow cytometry, and used immediately for experiments.

#### Generation, Titration, and Intratracheal Administration of Cre Lentivirus into KP Mice

We previously described modification of the HIV7 lentiviral vector to encode truncated human ROR1 (hROR1t) (UniProt: Q01973, aa 1-462) linked by a P2A ribosomal skip element to Cre recombinase (hROR1t-P2A-Cre).^5,42^ We modified the same vector to co-express hROR1t, tCD19 (UniProt: P25918, aa1-321) fused via a SSSSG linker to the MHC class I-restricted epitope derived from chicken ovalbumin (UniProt: P01012, aa249-271), and Cre recombinase, with each transgene linked by P2A and T2A ribosomal skip elements (hROR1t-P2A-tCD19-Ova-T2A-Cre). Lentivirus was produced by transient calcium phosphate transfection of the packaging cell line LentiX with the indicated HIV7 lentiviral vectors, psPAX2 (Addgene plasmid # 12260), and VSVg envelope (Addgene plasmid #8454). Viral supernatant was harvested 24, 48 and 72 hr after transfection, filtered through a 0.45-μm pore filter, and stored at 4°C for up to 1 week until ready for ultracentrifugation. Lentivirus was concentrated by mixing filtered lentivirus with 40% polyethylene glycol (PEG, Sigma) at a PEG to virus ratio of 1:3 for 12-24 hr at 4°C. The virus/PEG mixture was then centrifuged at 1500g for 45 min at 4°C, supernatant was aspirated, and the virus pellet was resuspended in 30 ml serum-free DMEM. Lentivirus was further concentrated in an Optima L-90K ultracentrifuge (Beckman Coulter) at 24,500 rpm for 90 min at 4°C. The final virus pellet was resuspended in 0.5 – 1 ml serum-free DMEM by vortexing for 1-3 hr at 4°C, aliquoted, and frozen at -80°C for long-term storage.

Cre lentivirus was titered using 3TZ “Green Go” cells. Briefly, 5×10^4^ 3TZ cells were plated in 1 ml of complete DMEM in 12-well plates. 5-6 hours later when cells were adherent, supernatant was aspirated and media was replaced with 0.5 ml complete DMEM containing serial dilutions of thawed Cre lentivirus (e.g. 10-fold serial dilutions from 1:10 to 1:10,000) and 8 μg/ml polybrene. After 24 hr, media was replaced with 1 ml complete DMEM and cells were passaged for 3-4 days before analysis by flow cytometry for GFP, hROR1, and/or tCD19 expression. Virus titer was calculated according to the following formula: Titer (pfu/μl) = [(5×10^4^) * (%GFP+ cells) / 100] / [Volume of virus added (μl)].

#### Tumor Models

2.5×10^4^ KP^ROR1^ tumor cells were injected intravenously via tail vein into 8-12 week old B6 mice to induce development of metastatic lung tumors. Mice were monitored for weight, body condition score, tumor burden, and survival where indicated. For subcutaneous tumor models, the right flank was shaved and one day later, 2x10^6^ KP^ROR1-Ova^ tumor cells were injected subcutaneously. Tumors were measured via digital calipers, and tumor volume was calculated using the formula V = ((length^2)*width)/2). Mice were euthanized when any tumor dimension reached 2 cm or when >80% of the tumor surface was ulcerated. For the autochthonous KP^ROR1^ model, lung tumors were induced in KP mice by intratracheal intubation and inhalation of 2x10^4^ pfu ROR1-P2A-Cre lentivirus or 5x10^4^ pfu ROR1-P2A-CD19-Ova-T2A-Cre lentivirus. Mice were monitored for weight, body condition score, tumor burden, and survival where indicated. Mice were sacrificed when they showed clinical signs of severe disease and/or >20% weight loss.

#### CAR-T Cell Treatment

KP^ROR1^ and KP^ROR1-Ova^ mice were scanned by MRI 10-11 weeks post-infection and one week before T cell infusion, and tumor burden was calculated for each mouse using ImageJ or Vivoquant software (Invicro). For subcutaneous KP^ROR1-Ova^ tumors, tumor volume was measured by digital calipers. Mice were distributed into treatment groups such that the average tumor burden was comparable across experimental groups prior to treatment. For all CAR-T cell infusion experiments, mice were injected intraperitoneally with 150 mg/kg cyclophosphamide and 6-24 hr later were injected intravenously by retro-orbital injection with 6×10^6^ live tCD19^+^ CAR-T cells (1:1 ratio of CD8:CD4 T cells, 3x10^6^ of each). Treatment was administered 10 days post-tumor injection for the subcutaneous KP^ROR1-Ova^ tumor model and 14-21 days post-tumor injection for KP^ROR1^ lung metastasis tumor model, when mice had visible tumor nodules in the lungs. For autochthonous KP^ROR1^ and KP^ROR1-Ova^ models, treatment was started 12-15 weeks post-infection with lentivirus and continued every 3 weeks.

#### NKTR-255 Treatment

NKTR-255 and vehicle dilution buffer were generously provided by Nektar Therapeutics (San Francisco, CA). Mice were injected intravenously with 0.33mg/kg NKTR-255 or vehicle beginning 1 day post Cy and CAR-T cell treatment and once every 7 days thereafter.

#### Anti-Thy1.2 Antibody Treatment

Tumor-bearing mice were treated intraperitoneally with 250ug anti-Thy1.2 antibody (Clone 30H12, BioXcell, Cat: BE0066) diluted in PBS as previously described.^43^ Briefly, anti-Thy1.2 or vehicle control were administered one day prior to administration of Cy + CAR-T cells (Day -1), the day after (Day 1), and every 7 days for the remainder of the experiment.

#### MRI

Mice were imaged on a 1-Tesla Bruker ICONÔ MRI scanner. Animals were anesthetized with 1–3% isoflurane via induction chamber, then maintained on a nose cone. A gradient echo flow compensated sequence using a repetition time of 592.4 ms, echo time of 7.0 ms and flip angle of 80 were used throughout the study. The slice thickness was 1 mm, and the number of slices was 15, which was sufficient to cover the entire lung. The acquisition matrix size was 128 x 178, the reconstructed matrix size was 128 x 256, and the field of view was 25 x 25 mm2. Motion artifacts were minimized by application of respiratory gating to all MRI studies. All animals were scanned by using the described settings and parameters. Tumor burden was analyzed using ImageJ or Vivoquant software (Invicro). Briefly, region of interest (ROI) tools were used to annotate individual lung tumor nodules in each mouse at each time point. Volume of each tumor nodule was calculated by multiplying the tumor area (mm^2^) by the slice thickness (1 mm). Percent change in tumor volume was calculated using the following formula: [(Vol. at Time_X­_) – (Vol. at Time_0­_)] / [Vol. at Time_0­_] * 100, where Time_0­_is one week prior to CAR-T cell treatment.

#### Preparation of Tissues for Flow Cytometry and RNAseq

Blood was collected from mice using submental bleeding and placed in BD Vacutainer EDTA Tubes. Blood volume was measured with a pipet, and red blood cells were lysed by two sequential incubations with 1mL of ACK lysis buffer (Gibco) for 5 minutes each, followed by washing with FACS buffer (PBS + 2% FBS).

For analysis of lung tumors, mice were injected with 400 ng PECy7-conjugated anti-CD45 antibody (clone 30-F11, BioLegend) intravenously via retro-orbital injection 5 min prior to euthanasia to distinguish vascular and non-vascular cells in the lung. Whole lungs were placed in GentleMACS C Tubes with 6 ml of digestion buffer (complete RPMI supplemented with 2 mg/ml collagenase IV (Worthington) and 80 U/ml DNAse I (Worthington)). Lungs were minced using the “m_impTumor_01_01” program on a GentleMACS Octo Dissociator and incubated in a 37°C shaker at 120 rpm for 30 min, then were further dissociated using the “m_Lung_02_01” program. Cells were filtered through a 100 μm filter, lysed with ACK lysing buffer (Gibco), and resuspended as single cell suspensions for downstream analysis. All flow cytometry analyses of tumor-bearing lungs were performed on live CD45-PECy7-cells. Subcutaneously implanted tumors were dissected from mice, minced in 3mL of complete RPMI, and transferred to GentleMACS C Tubes containing 3mL of 2x digestion buffer (complete RPMI + 1.96mg/mL collagenase Type I + 160U/mL DNAse I). Tumors were then digested as described above.

Lung tumor-draining mediastinal lymph nodes were collected in complete RPMI and transferred into digestion mix (serum-free RPMI + 0.385mg/mL Collagenase I + 1.408 mg/mL Collagenase IV + 80 U/mL DNAseI). Tissues were minced by hand, incubated for 15 minutes at 37C, pipet mixed with a P1000, then incubated for another 15 minutes, and pipet mixed again. Digestion was quenched with an equal volume of FACS buffer (PBS + 2% FBS). Cells were filtered through a 35um filter on a 5mL Falcon® Round-Bottom Tubes with Cell Strainer Cap.

#### Flow Cytometry

For flow cytometry panels using peptide/MHC tetramers, cells were first treated with 200uL of 50nM dasatanib (Axon 1392) diluted in FACS buffer for 30 min at 37C to maximize surface TCR expression, as previously described.^44^ Cells were then stained using either the Live/Dead Fixable Aqua Dead Cell stain kit (Invitrogen) at 1:200 in 200 uL of PBS per sample, or the Zombie UV Fixable Viability Kit (BioLegend) at 1:1000 in 100uL PBS per sample, and incubated for 20 min at 4C. For surface staining, cells were incubated with antibodies diluted in FACS buffer (PBS, 2% FBS) for 30 mins at 4°C, or at 37°C when using peptide/MHC tetramers. H-2Kb/SIINFEKL peptide/MHC tetramers were generated by the Fred Hutch Immune Monitoring Core and were added into surface stains at a 1:100 dilution. Recombinant Fc-hROR1 (Fred Hutchinson Cancer Center Protein Core) and anti-human IgG Fc secondary (Clone M1310G05, Biolegend 410710) were used to measure surface ROR1 CAR expression and were added to surface stains at 1:500 and 1:100 dilutions, respectively. Cells were then fixed and permeabilized with the eBioscience Foxp3 Transcription Factor Staining Kit (Thermo Fisher) using manufacturer’s instructions. For intracellular staining, cells were stained with antibodies targeting intracellular proteins diluted in wash buffer for 30 minutes at 4C. For intracellular cytokine staining, 1x10^6^ cells from digested tumor cell suspensions were stimulated with 50ng/ml PMA and 1 μg/ml ionomycin in 0.2 ml complete RPMI in 96-well U-bottomed plates (Costar) at 37°C, 5% CO2 for 6 hr. GolgiPlug (BD Biosciences) was added to all wells according to the manufacturer’s protocol at the beginning of co-culture. Cells were surface stained, fixed and permeabilized with BD Cytofix/Cytoperm buffer, and stained with antibodies targeting intracellular proteins diluted in wash buffer, as described above. All data were acquired on FACS Symphony or Canto II flow cytometers (BD Biosciences) and analyzed using FlowJo software (Treestar). Flow antibodies used in each panel are listed in Supplementary Table S1.

Absolute numbers of various immune cells in tissues were determined using Polybead polystyrene nonfluorescent microspheres (15 um, Polysciences, Inc.). Briefly, 10ul of the cell suspension to be counted was stained with 100uL of Live/Dead Fixable Aqua diluted 1:200 in PBS for 20 minutes at 4C in a 96-well plate. 50ul of a fixed concentration (C_B­_) of Polybeads (one drop of Polybeads per ml of PBS) was then added to each well, followed by addition of 100ul of 4% paraformaldehyde to fix cells. Without washing, the samples were acquired on a FACS Celesta (BD) and were quantified using the appropriate gates, with SSC set to log-scale to visualize both cells and beads. Beads and lymphocytes were identified by their distinct forward- and side-scatter characteristics. The ratio of lymphocyte gate events (n_L­_) to bead gate events (n_B­_) was determined and used to calculate the concentration (C) of the original cell suspension as follows: C = (n_L­_/ n_B­_) * C_B­_

#### RNA Sequencing

Cryopreserved lung cell suspensions were thawed and stained with Live/Dead Fixable Aqua Dye (Invitrogen) and surface markers in FACS Buffer. One thousand live CD45-PECy7^-^CD8^+^tCD19^+^ CAR-T cells were sorted directly into reaction buffer from the SMART-Seq mRNA Kit (Takara), and reverse transcription was performed followed by PCR amplification to generate full-length amplified cDNA. Sequencing libraries were constructed using the NexteraXT DNA sample preparation kit with unique dual indexes (Illumina) to generate Illumina-compatible barcoded libraries. Libraries were pooled and quantified using a Qubit® Fluorometer (Life Technologies). Sequencing of pooled libraries was carried out on a NextSeq 2000 sequencer using an XLEAP-SBS flowcell (Illumina) with paired-end 59-base reads (Illumina) with a target depth of 5 million reads per sample. Base calls were processed to FASTQs on BaseSpace (Illumina). A base call quality-trimming step was applied to remove low-confidence base calls from the ends of reads using fastp (0.23.1). The FASTQs were aligned to the Ensemble mouse genome assembly version GRCm38.91 using STAR (2.7.11a), and gene counts were generated using htseq-count (2.0.2). QC and metrics analysis was performed using fastQC (0.12.1) and the Picard family of tools (3.1.0). Filtered and normalized gene counts were generated from the raw counts by trimmed-mean of M values (TMM) normalization and filtered for genes that are expressed with at least 1 count per million total reads in at least 10% of the total number of libraries. Differential expression analysis was performed on normalized gene counts using DESeq2 in R using R Studio version 2025.05.1+513. Log fold change shrinkage was performed using apeglm. Significant results were determined through |Log2FC| > 1 and the FDR adjusted p-value < 0.05. Volcano plots were made with EnhancedVolcano package. Heatmaps were made with pheatmap package.

#### Immunohistochemistry (IHC)

Tumor-bearing lungs were harvested from mice and directly placed into 10% neutral buffered formalin (NBF) for at least 3 days. Formalin-fixed paraffin-embedded tissues were sectioned at 4-5µm onto positively charged slides and baked for a minimum of 30 minutes at 65°C / 60 minutes at 60°C. The slides were then dewaxed and stained on a Leica BOND RX autostainer (Leica, Buffalo Grove, IL) using Leica Bond reagents for deparaffinization (Bond Dewax Solution, AR9222).

Antigen retrieval was conducted at 100°C for 20 minutes with Bond Epitope Retrieval Solution 2 (Leica, AR9640) for slides stained with ROR1 and GFP or Bond Epitope Retrieval Solution 1 (Leica, AR9961) for slides stained CD8a. Endogenous peroxidase was blocked using 3% H₂O₂ (5 minutes), followed by protein blocking with TCT buffer (0.05M Tris, 0.15M NaCl, 0.25% Casein, 0.1% Tween 20, 0.05% ProClin300, pH 7.6) for 10 minutes. ROR1 (D6T8C; Cell Signaling Technology, 16540 at a 1:500 dilution) and GFP (rabbit polyclonal; Invitrogen, A11122 at a 1:800 dilution) were applied to respective slides for 60 minutes at room temperature. For detection, slides were then incubated with PowerVision Poly-HRP anti-rabbit (Leica PV6119) for 12 minutes, respectively. Mixed Refine DAB (Leica DS9800) was applied to all slides for 10 minutes, followed by counterstaining with Refine Hematoxylin (Leica DS9800) for 4 minutes. Slides were then dehydrated, cleared, and cover-slipped with permanent mounting media.

Percent ROR1^+^ cells within tumors was quantified using HALO software (Indica Labs). A classifier based on Random Forest was generated. Percent GFP^+^ cells in tumors were quantified using QuPath, where tumor borders were drawn manually and positive cell detection was used to identify GFP^+^ and GFP^-^ cells in the tumors.

#### Quantification and Statistical Analysis

Statistical significance was determined by two-way unpaired Student’s t-test, one-way ANOVA with Tukey’s post-test, multiple t-test with Holm-Šidák correction, two-way ANOVA with Holm-Šidák correction, or log-rank Mantel-Cox test, as indicated in figure legends using Prism software (Graphpad). Statistical significance was established at the levels of *, p<0.05; **, p<0.005; ***, p<0.0005; ****, p<0.0001.

## Results

### NKTR-255 significantly increases CAR-T cell accumulation in vivo

To investigate the effects of NKTR-255 on both CAR-T cells and endogenous immune cells, we adapted the *Kras^LSL-G12D^*^/+^;*p53^fl/fl^*(**KP**) autochthonous model of NSCLC to express truncated human ROR1 (**KP^ROR1^**).^5,38,39,45^ We engineered primary murine CD8^+^ and CD4^+^ T cells to co-express a ROR1-targeted CAR containing 4-1BB and CD3ς signaling domains and truncated CD19 (**tCD19**) as a transduction marker (Fig. 1A).^5^ We previously showed that this model mimics the poor persistence and dysfunction of ROR1 CAR-T cells observed in NSCLC and TNBC patients in a phase 1 trial at our Center, and that serial infusion of CD8^+^ and CD4^+^ ROR1 CAR-T cells was unable to confer a significant survival benefit.^4,5,42^ To determine whether NKTR-255 could overcome the poor accumulation and persistence of ROR1 CAR-T cells, we treated tumor-bearing KP^ROR1^ mice every three weeks with cyclophosphamide (**Cy**) to induce lymphodepletion, infused CD8^+^ and CD4^+^ ROR1 CAR-T cells, and administered NKTR-255 or vehicle one day later and weekly thereafter (Fig. 1B). Treatment was well tolerated, with no significant change in weight loss or body condition score observed (Fig. S1A, S1B), though mice treated with CAR-T cells and NKTR-255 showed an increase in spleen mass by 8 weeks after the first CAR-T infusion, a common effect observed with modified IL-15 drugs (Fig. S1C).^46,47^ While CAR-T cells accumulated poorly in the blood of vehicle-treated mice, NKTR-255 treatment dramatically increased both CD8^+^ and CD4^+^ CAR-T cell frequency (Fig. 1C, 1D). These effects were more pronounced amongst CD8^+^ CAR-T cells, which accumulated to ∼50-fold higher levels with NKTR-255, compared to CD4^+^ CAR-T cells, which increased by ∼3-fold, despite being infused at equal numbers.

**Figure 1.**
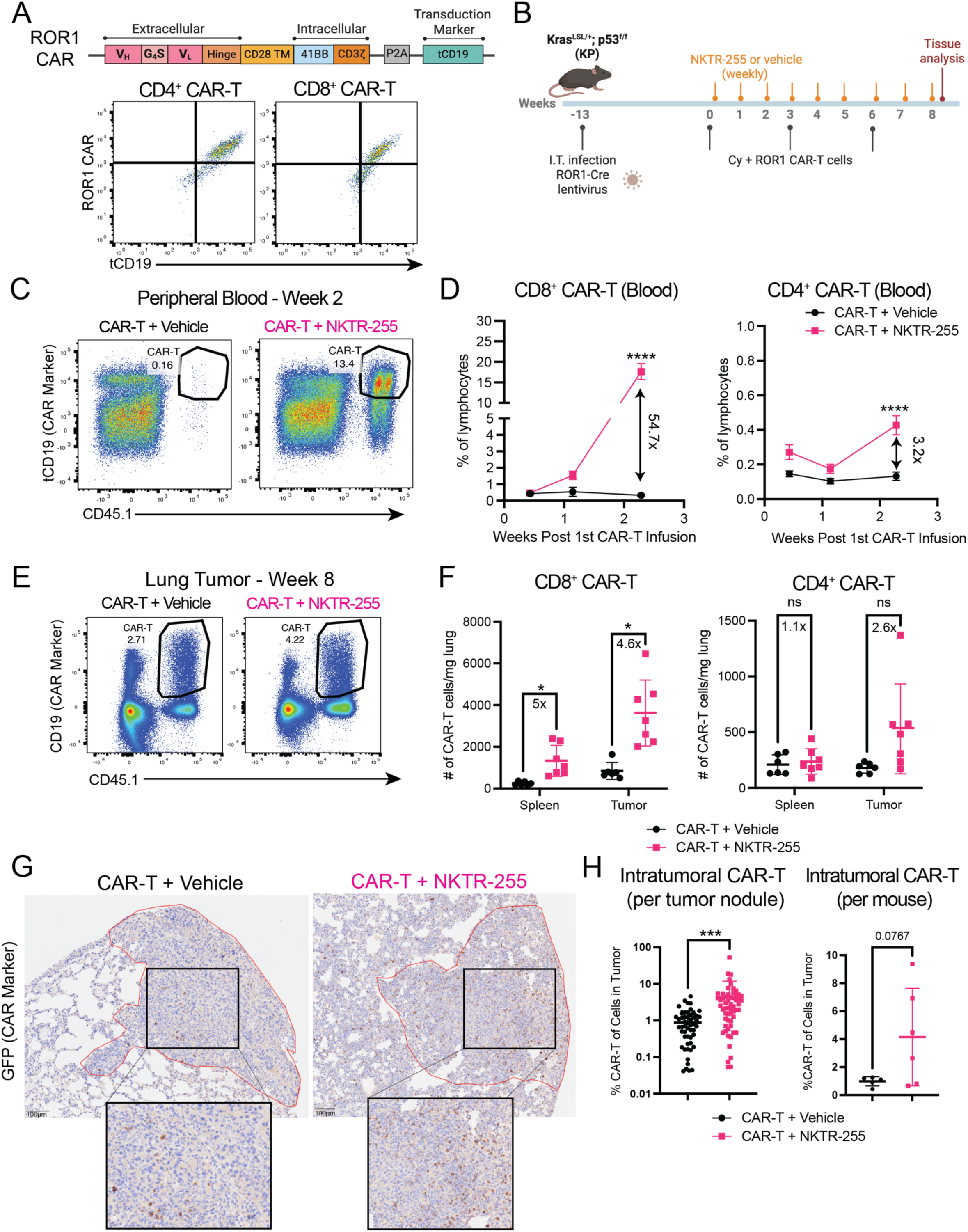
NKTR-255 significantly expands CAR-T cells *in vivo*. A) Top: Schematic of CAR construct. TM = transmembrane. tCD19 = truncated CD19. Bottom: representative flow plots showing CAR and tCD19 expression in CD8^+^ and CD4^+^ ROR1 CAR-T cell infusion products. B) Experimental schematic. I.T. = intratracheal. Cy = cyclophosphamide. C) Representative flow plots showing CD45.1^+^CD19^+^ CAR-T cell frequency of total lymphocytes in blood 2 weeks post-CAR-T infusion. D) CD45.1^+^CD8^+^CD19^+^ and CD45.1^+^CD4^+^CD19^+^ CAR-T frequency of total lymphocytes in blood. Mean +/- SEM. Two-way ANOVA with Holm-Šidák correction. N=13-14 mice per group. E) Representative flow plots showing CD45.1^+^CD19^+^ CAR-T cell frequency of total lymphocytes in lung tumors 2 weeks post-CAR-T infusion (8 weeks post-1^st^ CAR-T infusion). F) Absolute number of CD45.1^+^CD8^+^CD19^+^ and CD45.1^+^CD4^+^CD19^+^ CAR-T cells per mg of tissue 2 weeks post-CAR-T infusion (8 weeks post-1^st^ CAR-T infusion). Mean +/- SD. Multiple t-test with Holm-Šidák correction. N=6-7 mice per group. G) Representative immunohistochemistry images of GFP^+^ CAR-T cells in lung tumor nodules 10 days post-CAR-T infusion. H) Frequency of CAR-T cells as a fraction of total cells in individual tumor nodules (left) or as a fraction of total tumor area (right). Mean +/- SD. Two-way unpaired Student’s t-test. N=58-59 tumor nodules per group (left) from N=5-6 mice per group (right).

To determine if NKTR-255 similarly expanded ROR1 CAR-T cells in tissues, we next examined CAR-T cell accumulation in the spleen and tumor-bearing lung. To discriminate cells in lung tissue/tumors from circulating cells in lung vasculature, we used a previously described *in vivo* labeling method where tumor-bearing mice are injected intravenously with a PECy7-conjugated anti-CD45 antibody just prior to euthanasia and performed subsequent analyses on live CD45 PECy7^-^ non-vascular cells in the lung.^48–50^ Similar to the blood, NKTR-255 treatment increased the frequency of CD8^+^ CAR-T cells in both spleens and lung tissue, though the accumulation of CD8^+^ CAR-T cells was greatest in lungs (Fig. 1E, 1F). By contrast, NKTR-255 treatment did not significantly increase CD4^+^ CAR-T cells in the spleen and induced only a slight, but insignificant, increase within lungs, consistent with the more pronounced effects of NKTR-255 on CD8^+^ T cells in the blood (Fig. 1F). To validate whether NKTR-255 increased CAR-T cells within lung tumor nodules, we modified the CAR vector to co-express tCD19 fused to GFP as a transduction marker and used immunohistochemistry (**IHC**) to visualize GFP^+^ CAR-T infiltration within KP^ROR1^ tumors. Consistent with our flow cytometry data, we found NKTR-255 significantly increased GFP^+^ CAR-T cell density within lung tumors (Fig. 1G-1H). Thus, NKTR-255 dramatically increases CAR-T cell accumulation in both tumors and the periphery.

### NKTR-255 reduces exhaustion and enhances the pro-inflammatory function of CAR-T cells within lung tumors

We previously showed that poor efficacy of ROR1 CAR-T cells was correlated with rapid upregulation of inhibitory receptors and decreased capacity to produce pro-inflammatory cytokines in both KP^ROR1^ mice and NSCLC patients.^5^ While IL-15 is known to enhance T cell survival, proliferation and stemness,^51^ it is not clear how it impacts the development of CAR-T exhaustion and dysfunction in an immunosuppressive tumor microenvironment. As the effects of NKTR-255 were most pronounced on CD8^+^ CAR-T cells, we focused our analysis on CD8^+^ CAR-T cells, which comprised the vast majority of CAR-T cells within lung tumors. While lung tumor-infiltrating CAR-T cells in vehicle-treated mice expressed uniformly high levels of PD-1, CAR-T cells in NKTR-255-treated mice showed significantly lower PD-1 expression, with a substantial fraction of cells remaining PD-1^-^ (Fig. 2A). This effect appeared to be driven by a preferential increase in the number of PD-1^-^CAR-T cells and minimal change in the number of PD-1^+^ CAR-T cells (Fig. 2B), suggesting NKTR-255 preferentially expands less differentiated CAR-T cells.^30,52^ Although PD-1^+^ CAR-T cells were not increased by NKTR-255, they expressed PD-1 at significantly lower levels than PD-1^+^ CAR-T cells in vehicle-treated mice (Fig. 2C). As higher PD-1 levels correlate with increased T cell exhaustion/dysfunction,^53–56^ these data suggest that NKTR-255 may also favorably impact the differentiation of PD-1^+^ CAR-T cells in addition to its effects on expanding less differentiated PD-1^-^ CAR-T cells. Consistent with this hypothesis, PD-1^+^ CAR-T cells showed significantly reduced co-expression of the inhibitory receptor LAG-3 and the master transcription factor of T cell exhaustion, Tox (Fig. 2D, 2E), suggesting NKTR-255 promotes expansion of less terminally exhausted CAR-T cells.^57–62^

**Figure 2.**
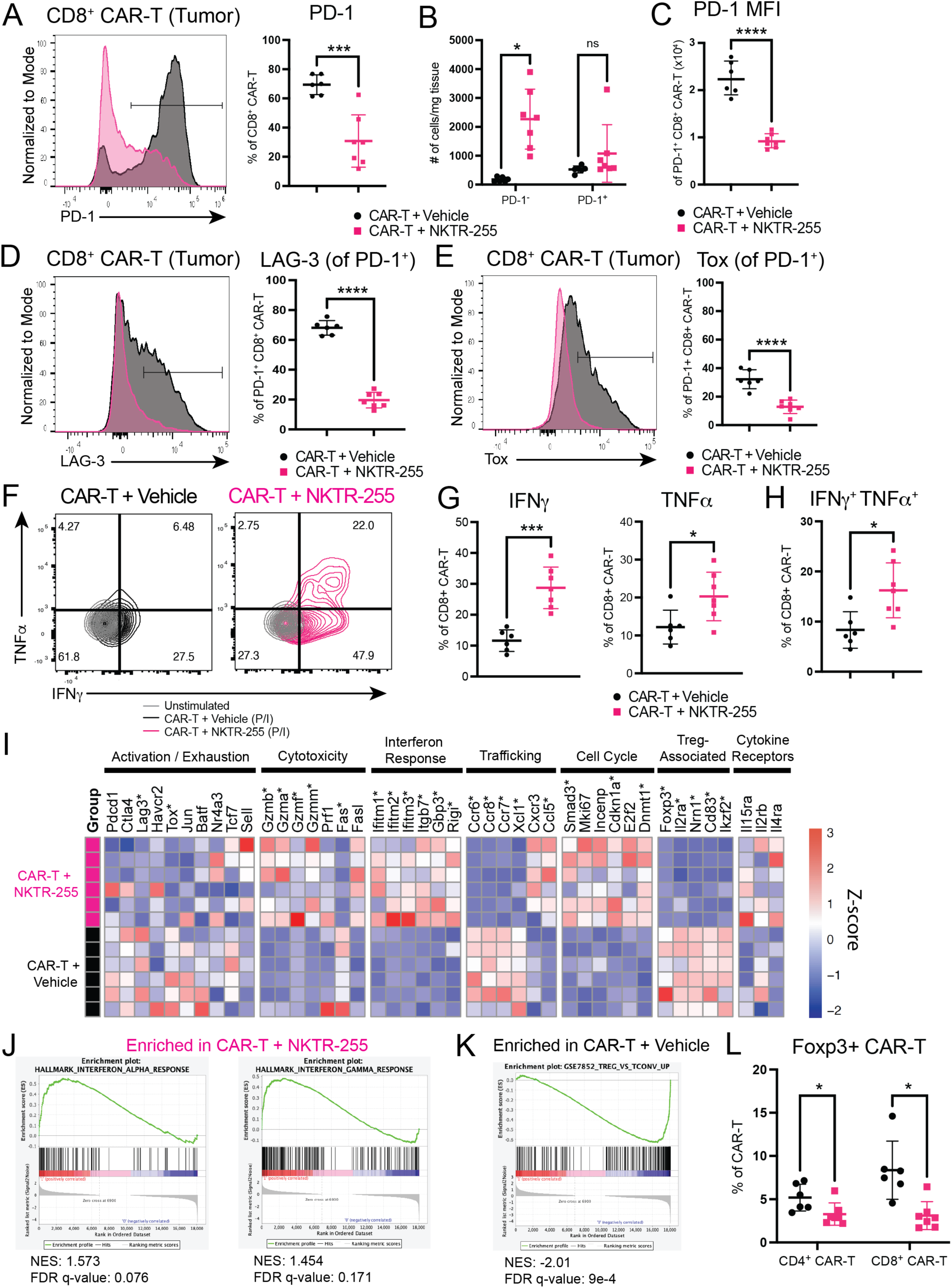
NKTR-255 enhances the pro-inflammatory function of CAR-T cells within tumors. A) PD-1 expression on CD8^+^CD19^+^ CAR-T cells in KP^ROR1^ lung tumors 2 weeks post-CAR-T infusion. Mean +/- SD. Two-way unpaired Student’s t-test. N=6-7 mice per group. B) Absolute number of PD-1^+^ and PD-1^-^ CD8^+^CD19^+^ CAR-T cells in KP^ROR1^ lung tumors 2 weeks post-CAR-T infusion. Mean +/- SD. Multiple unpaired Student’s t-test with Šidák correction. N=6-7 mice per group. C) Geometric mean fluorescence intensity (**MFI**) of PD-1 on PD-1^+^CD8^+^CD19^+^ CAR-T cells in KP^ROR1^ lung tumors 2 weeks post-CAR-T infusion. Mean +/- SD. Two-way unpaired Student’s t-test. N=6-7 mice per group. D, E) LAG-3 and Tox expression in PD-1^+^CD8^+^CD19^+^ CAR-T cells in KP^ROR1^ lung tumors 2 weeks post-CAR-T infusion. Mean +/- SD/ Two-way unpaired Student’s t-test. N=6-7 mice per group. F, G) Representative flow plots (F) and summary (G) of IFNγ and TNFα production by CD8^+^CD19^+^ CAR-T cells in KP^ROR1^ lung tumors 2 weeks post-CAR-T infusion after *ex vivo* restimulation with PMA/ionomycin. Mean +/- SD. Two-way unpaired Student’s t-test. N=6-7 mice per group. H) Frequency of CD8^+^CD19^+^ CAR-T cells in KP^ROR1^ lung tumors co-producing IFNγ and TNFα 2 weeks post-CAR-T infusion. Mean +/- SD. Two-way unpaired Student’s t-test. N=6-7 mice per group. I) Heatmap of z-score normalized counts of indicated genes in CD8^+^CD19^+^ CAR-T cells sorted from KP^ROR1^ lung tumors of vehicle- or NKTR-255-treated mice 2 weeks post-CAR-T infusion. * indicates genes differentially expressed between vehicle and NKTR-255 treatment (|log2FC|> 1 and adjusted p-value < 0.05). N=6 mice per group. J, K) GSEA plots of genesets significantly enriched in CD8^+^CD19^+^ CAR-T cells sorted from tumors of NKTR-255-treated (J) or from vehicle-treated KP^ROR1^ mice (K). L) Foxp3 expression in CD8^+^CD19^+^ and CD4^+^CD19^+^ CAR-T cells in KP^ROR1^ lung tumors 2 weeks post-CAR-T infusion. Mean +/- SD. Multiple unpaired Student’s t-test with Holm-Šidák correction. N=6-7 mice per group.

To determine whether NKTR-255 could rescue the function of tumor-infiltrating CAR-T cells, we restimulated CAR-T cells from lung tumors *ex vivo* and measured their capacity to produce pro-inflammatory cytokines. Whereas CAR-T cells from tumors of both vehicle- and NKTR-255-treated mice lost the ability to produce IL-2 (Fig. S2A), a greater proportion of CAR-T cells from tumors of NKTR-255-treated mice retained the capacity to produce IFNγ and TNF𝑎 (Fig. 2F, 2G). In particular, NKTR-255 promoted an increase in polyfunctional CAR-T cells that co-produced both IFN𝛾 and TNF𝑎, a characteristic associated with enhanced activity *in vivo* (Fig. 2H).^63,64^ These data suggest NKTR-255 significantly reduces exhaustion and preserves function of CAR-T cells in the lung tumor microenvironment.

To more comprehensively determine how NKTR-255 impacted the phenotype of ROR1 CAR-T cells within lung tumors, we sorted CAR-T cells from lung tumors and used bulk RNA sequencing to compare their transcriptional profiles. Differential expression analysis identified 240 genes significantly upregulated and 345 genes significantly downregulated in CAR-T cells from NKTR-255-treated mice compared to CAR-T cells from vehicle-treated mice (Fig. S2B). Consistent with our flow cytometry data, CAR-T cells from NKTR-255-treated mice expressed lower levels of the exhaustion-associated transcripts *Tox* and *Lag3*. They also expressed higher levels of multiple granzymes (*Gzma*, *Gzmb*, *Gzmf*, *Gzmm*), and higher levels of cell cycle genes, suggesting increased cytotoxicity, effector function, and proliferation (Fig. 2I). Additionally, gene-set enrichment analysis (**GSEA**) showed that genesets associated with IFNγ and IFNα response were significantly enriched among CAR-T cells from NKTR-255-treated mice (Fig. 2I, 2J), consistent with their increased IFN𝛾 production that may promote a more inflammatory tumor microenvironment. Interestingly, several studies have observed that a subset of tumor-infiltrating CAR-T cells upregulate Foxp3, the master transcriptional regulator of regulatory T cells (**Tregs**), and that this may drive a suppressive transcriptional program that may contribute to T cell dysfunction in the TME.^65–68^ Indeed, gene sets associated with Foxp3^+^ regulatory T cells (**Tregs**) were significantly enriched among CAR-T cells from vehicle-treated mice (Fig. 2K), and Foxp3 protein levels were detectable in both tumor-infiltrating CD8^+^ and CD4^+^ CAR-T cells of vehicle-treated mice, but were significantly reduced with NKTR-255 treatment (Fig. 2L). Thus, NKTR-255 treatment significantly reduces CAR-T cell exhaustion, enhances their pro-inflammatory function, and may protect them from upregulation of immunoregulatory gene signatures in lung tumors.

### NKTR-255 and CAR-T cell combination treatment enhances endogenous tumor-specific T cell responses in lung tumors

In addition to its effects on CAR-T cells, an attractive aspect of NKTR-255 therapy is its potential to also activate endogenous tumor-specific T cells that could functionally contribute to tumor control. However, a standard component of all CAR-T cell therapy regimens is the administration of lymphodepleting chemotherapy prior to CAR-T cell infusion, which may blunt endogenous tumor-specific T cell responses.^69,70^ Thus, it is unclear whether NKTR-255 is able to effectively boost endogenous tumor-specific T cells in the context of lymphodepleting chemotherapy and CAR-T therapy, and how these effects compare to CAR-T or NKTR-255 monotherapy alone.

To evaluate how CAR-T and/or NKTR-255 therapy impact endogenous tumor-specific T cells, we adapted the KP^ROR1^ model to co-express the MHC class I-restricted peptide SIINFEKL derived from the model neoantigen ovalbumin (**KP^ROR1-Ova^**), enabling us to use peptide/MHC tetramers to track endogenous H-2K^b^/SIINFEKL-specific CD8^+^ T cells (**Endog. CD8^+^Tet^+^**). Previous work has shown that expression of SIINFEKL in KP tumors induces activation of Endog. CD8^+^Tet^+^ that delay tumor progression, but that these cells ultimately become dysfunctional and mediate incomplete tumor control.^38^ We modified the lentivirus used to induce KP tumors to express hROR1t, truncated CD19 (**tCD19**) fused to the SIINFEKL peptide, and Cre recombinase, with transgenes linked by P2A and T2A skip elements, and we confirmed co-expression of hROR1t and tCD19 in lentivirus-infected 3TZ cells (Fig. 3A).

**Figure 3.**
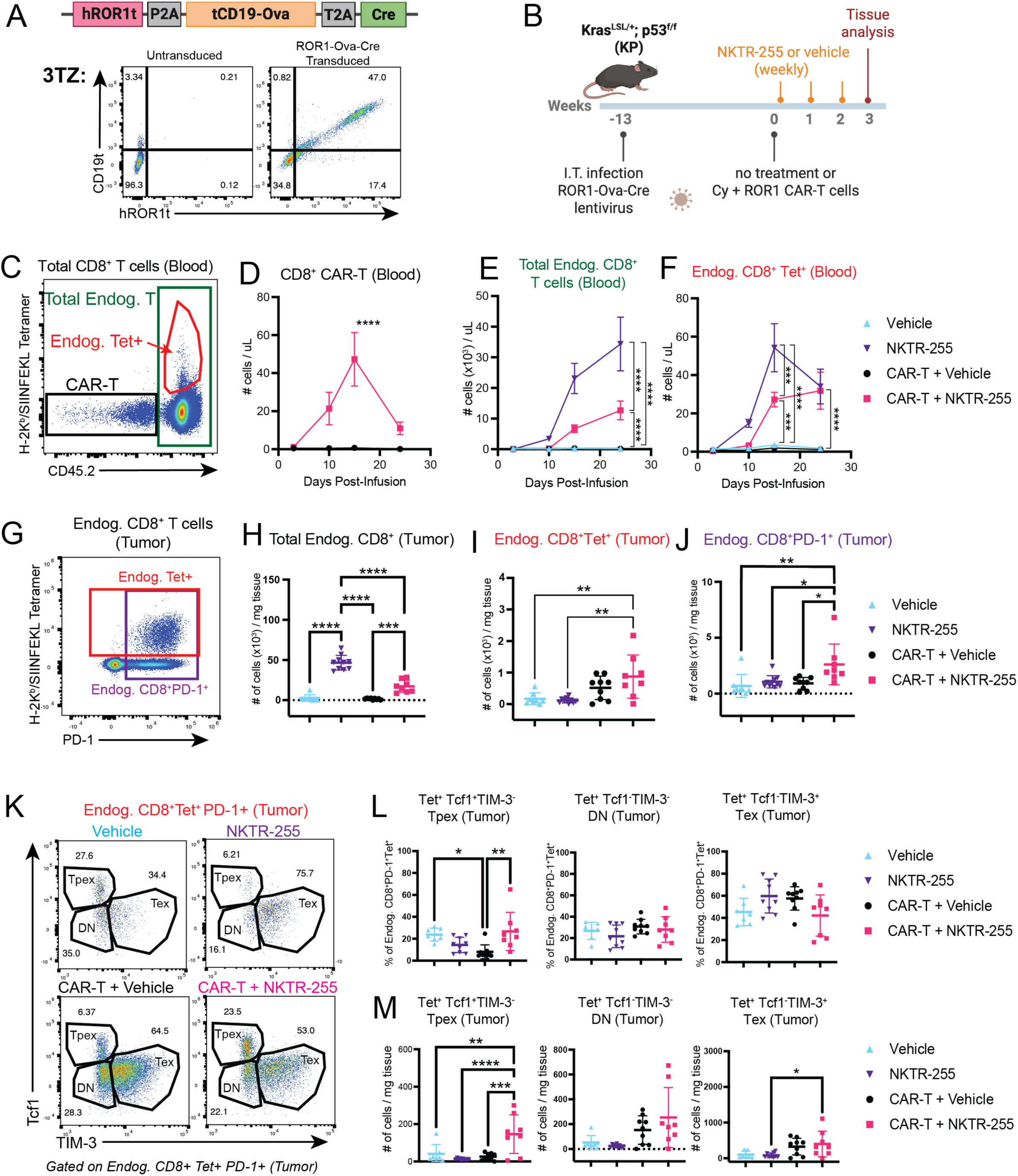
NKTR-255 and CAR-T cell combination therapy increases endogenous stem-like tumor-specific T cells within tumors. A) Top: schematic of ROR1-Ova-Cre lentivirus constructs used to induce ROR1^+^Ova^+^ lung tumors. hROR1t = truncated human ROR1. tCD19 = truncated murine CD19. Ova = ovalbumin. Bottom: co-expression of hROR1t and CD19t in 3TZ cells transduced with ROR1-Ova-Cre lentivirus. B) Experimental schematic. I.T. = intratracheal. Cy = cyclophosphamide. C) Gating strategy for various CD8^+^ subsets in blood of KP^ROR1-Ova^ mice. D-F) Number of CD8^+^CD19^+^ CAR-T cells (D), endogenous CD45.2^+^CD8^+^ T cells (E), and endogenous CD45.2^+^ H-2K^b^/SIINFEKL tetramer^+^ CD8^+^ T cells (**Endog. CD8^+^Tet^+^**) (F) in blood of KP^ROR1-Ova^ mice treated as indicated. Mean +/- SD. Two-way ANOVA with Holm-Šidák correction. N=6-7 mice per group. G) Gating strategy for various CD8^+^ subsets in KP^ROR1-Ova^ lung tumors. H-J) Number of total endogenous CD45.2^+^CD8^+^ T cells (H), endogenous CD45.2^+^ H-2K^b^/SIINFEKL tetramer^+^ CD8^+^ T cells (**Endog. CD8^+^Tet^+^**) (I), and endogenous CD45.2^+^CD8^+^PD-1^+^ T cells (J) in KP^ROR1-Ova^ lung tumors 3 weeks post-CAR-T infusion. Mean +/- SD. One-way ANOVA with Tukey’s post-test. N=8-10 mice per group. K) Representative flow cytometry plots of Tcf1 and Tim3 expression in endogenous CD45.2^+^ PD-1^+^ H-2K^b^/SIINFEKL tetramer^+^ CD8^+^ T cells in KP^ROR1-Ova^ lung tumors. L, M) Frequency (L) and number (M) of Tcf1^+^Tim3^-^ Tpex, Tcf1^-^Tim3^-^ double negative (DN) and Tcf1^-^Tim3^+^ Tex in KP^ROR1-Ova^ lung tumors 3 weeks post-CAR-T infusion. L-M: Mean +/- SD. One-way ANOVA with Tukey’s post-test. N=8-10 mice per group.

We treated KP^ROR1-Ova^ tumor-bearing mice with Cy and ROR1 CAR-T cells and administered either vehicle or NKTR-255, as before. To examine the effects of NKTR-255 monotherapy in the absence of lymphodepleting chemotherapy and CAR-T cells, we also treated subsets of KP^ROR1-Ova^ tumor-bearing mice with either vehicle or NKTR-255 alone (no Cy or CAR-T cells) (Fig. 3B). We first confirmed NKTR-255 treatment increased CD8^+^ CAR-T cell numbers in the blood of KP^ROR1-Ova^ mice, similar to our results in the KP^ROR1^ model (Fig. 3C, 3D). NKTR-255 monotherapy and combination therapy with CAR-T cells also promoted a significant increase in total endogenous CD8^+^ T cells in the blood, but the increase was significantly higher in mice treated with NKTR-255 monotherapy (Fig. 3E), presumably because pre-conditioning with Cy prior to CAR-T infusion reduced numbers of endogenous CD8^+^ T cells compared to mice treated with NKTR-255 alone. Interestingly, NKTR-255 monotherapy initially expanded Endog. CD8^+^Tet^+^ in the blood more effectively than CAR-T and NKTR-255 combination therapy; however, numbers of Endog. CD8^+^Tet^+^ in the blood of CAR-T and NKTR-255-treated mice eventually recovered by 3 weeks post-treatment to levels comparable to mice treated with NKTR-255 alone (Fig. 3F). These data suggest that NKTR-255 can promote systemic expansion of endogenous tumor-specific T cells both as a monotherapy, as well as in the context of Cy + CAR-T therapy, though with delayed kinetics.

We next examined how NKTR-255 monotherapy or combination therapy with CAR-T cells affected endogenous T cells within lung tumors. We analyzed lungs 3 weeks post-CAR-T infusion, when CAR-T + NKTR-255 combination treatment showed the greatest increase in Endog. CD8^+^Tet^+^ cells in the blood. In lung tumors, NKTR-255 treatment alone increased total endogenous CD8 T cell numbers to a greater extent than in CAR-T + NKTR-255 treated mice, mirroring what we observed in the blood (Fig. 3G, 3H). By contrast, we observed distinct effects of NKTR-255 on endogenous tumor-specific T cells within lung tumors: Endog. CD8^+^Tet^+^ T cells were not significantly increased in lung tumors following NKTR-255 monotherapy, despite showing an increase in the blood at this time point. Instead, Endog. CD8^+^Tet^+^ cells in tumors showed the most significant increase in response to CAR-T and NKTR-255 combination therapy, though their numbers were only slightly and insignificantly greater than CAR-T treatment alone (Fig. 3I). We observed a similar effect of NKTR-255 on endogenous CD8^+^PD-1^+^ T cells, which identifies antigen-experienced T cells within tumors and enriches for T cells reactive to tumor antigens in addition to H-2K^b^/SIINFEKL.^71,72^ Again, NKTR-255 or CAR-T monotherapy induced only a modest but insignificant increase in endogenous CD8^+^PD-1^+^ T cells within tumors, while CAR-T + NKTR-255 combination therapy induced the greatest and most significant increase in endogenous CD8^+^PD-1⁺ T cell numbers. These data suggest that NKTR-255 monotherapy is capable of expanding endogenous tumor-specific T cells in the blood but is unable to significantly increase these cells within tumors; by contrast, CAR-T and NKTR-255 combination therapy shows the greatest ability to increase tumor-reactive T cells within tumors, greater than either monotherapy alone.

In addition to promoting T cell accumulation, IL-15 has been shown to affect T cell differentiation and exhaustion, in particular by promoting self-renewal and proliferation of stem-like Tpex, which play a critical role in sustaining anti-tumor responses.^30^ Thus, we next measured how NKTR-255 and/or CAR-T therapy affected the phenotype of Endog. CD8^+^Tet^+^ in lung tumors. To determine how treatment impacted the differentiation of Endog. CD8^+^Tet^+^PD-1^+^ T cells into Tpex or into terminally exhausted subsets, we used Tcf1 and TIM-3 to define the following subsets of PD-1^+^ exhausted T cells within lung tumors: Tcf1^+^TIM-3^-^ defines Tpex cells, Tcf1^-^TIM-3^-^ double negative (**DN**) defines transitory exhausted cells, and Tcf1^-^TIM-3^+^ defines terminally exhausted cells (**Tex**).^73–75^ NKTR-255 monotherapy did not significantly impact the phenotype of Endog. CD8^+^Tet^+^ cells, suggesting that NKTR-255 alone has minimal impact on both the number and differentiation of endogenous tumor-specific cells (Fig. 3K, 3L, 3M). CAR-T monotherapy induced a significant reduction in the proportion of Tpex and increased the proportion of Tcf1^-^ DN and Tex subsets of Endog. CD8^+^Tet^+^ cells, suggesting that while CAR-T monotherapy can increase the number of tumor-specific cells in tumors, it promotes their terminal exhaustion. In contrast, CAR-T and NKTR-255 combination therapy significantly increased the proportion and number of Tpex among Endog. CD8^+^Tet^+^ T cells in tumors compared to either treatment alone, suggesting that combination therapy can uniquely increase stem-like tumor-specific cells within tumors.

### NKTR-255 and CAR-T cell combination therapy promotes increased DC activation in lung tumors

The ability of CAR-T cells to promote endogenous T cell activation and epitope spreading has been shown to be driven in part by increased activation of conventional dendritic cells (**cDCs**), which cross-present tumor-derived antigen to endogenous CD8^+^ T cells.^18,19,76^ Thus, we next examined how CAR-T and/or NKTR- 255 therapy affected cDC numbers and phenotype in tumors. We examined both cDC1 (Xcr1^+^CD11b^-^) and cDC2 (Xcr1^-^CD11b^+^) subsets of cDCs, as both are involved in regulating endogenous T cells in tumors (Fig. S3A).^77^ Neither cDC1 nor cDC2 numbers were significantly changed in tumors across any treatment group, though there was a trend towards increased cDC1 and cDC2 numbers in tumors of mice treated with NKTR-255, either as a monotherapy or in combination with CAR-T therapy (Fig. 4A). To examine the phenotype of tumor-infiltrating DCs, we measured expression of the activation and maturation markers CD40 and CD86, which have been shown to be upregulated in response to IL-15 *in vitro* as well as following adoptive T cell therapy.^19,76,78^ In tumors, cDC1s showed increased CD86 expression and slightly elevated CD40 expression after CAR-T + NKTR-255 combination therapy compared to either treatment alone (Fig. 4B). Conversely, cDC2s showed significantly increased CD40 and minimal changes to CD86 following CAR-T and NKTR-255 combination therapy (Fig. 4C). Co-expression of CD40 and CD86 mark “activated” DCs that correlate with increased endogenous tumor-specific T cell responses. Indeed, CAR-T and NKTR-255 combination therapy synergistically increased numbers of CD40^+^CD86^+^ activated cDC1 and cDC2 within tumors compared to either monotherapy (Fig. 4D).

**Figure 4.**
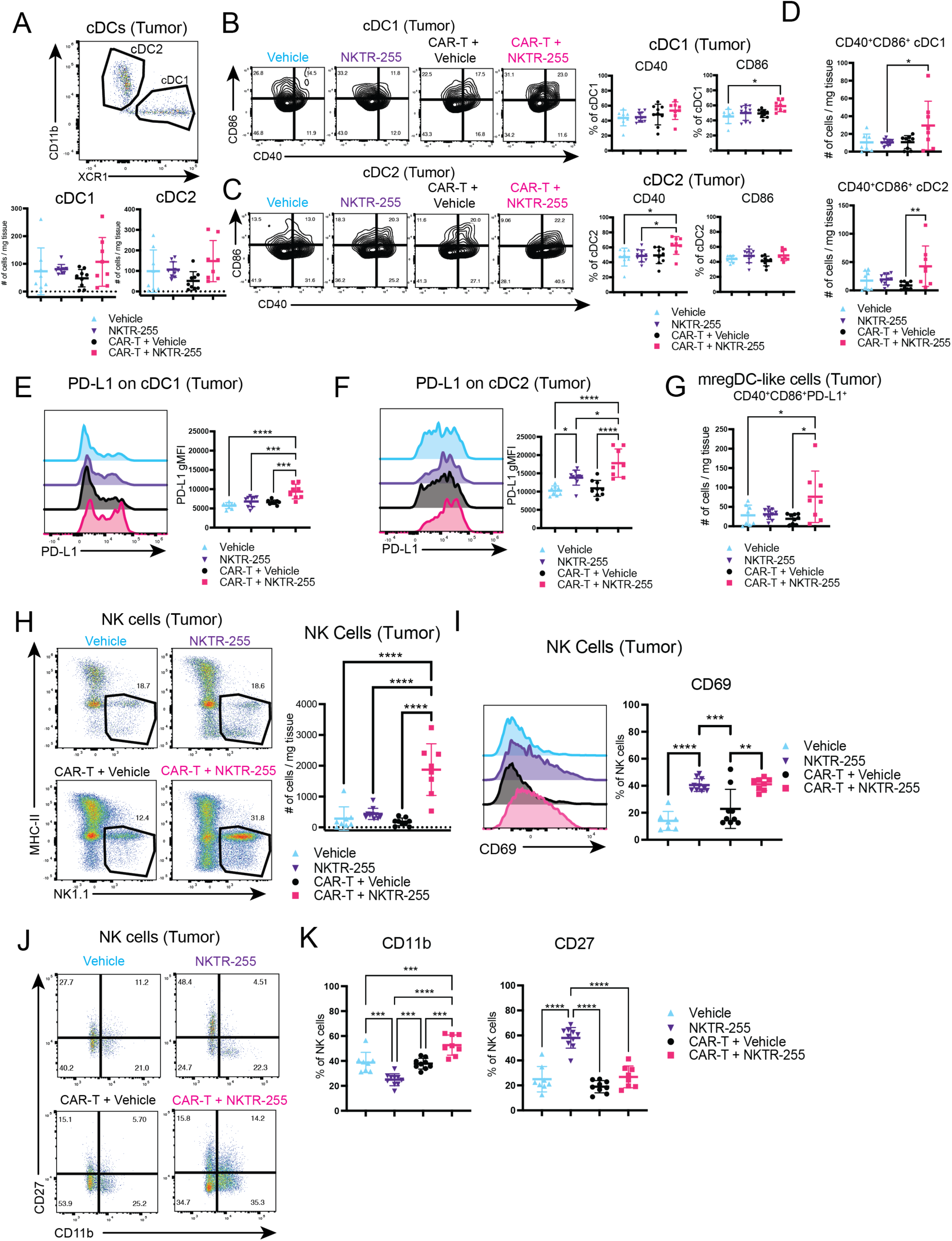
NKTR-255 and CAR-T cell combination therapy increases DC activation and cytotoxic NK cells in tumors. A) Top: cDC1 and cDC2 gating strategy. Bottom: number of cDC1 and cDC2 in KP^ROR1-Ova^ lung tumors 3 weeks post-CAR-T infusion. B, C) Representative flow plots (left) and frequency (right) of CD40 and CD86 expression on cDC1 (B) and cDC2 (C) in KP^ROR1-Ova^ lung tumors 3 weeks post-CAR-T infusion. D) Number of CD40^+^CD86^+^ cDC1 and cDC2 in KP^ROR1-Ova^ lung tumors 3 weeks post-CAR-T infusion. E, F) PD-L1 expression on cDC1 (E) and cDC2 (F) in KP^ROR1-Ova^ lung tumors 3 weeks post-CAR-T infusion. G) Number of CD40^+^CD86^+^PD-L1^+^ “mreg-like” DCs in KP^ROR1-Ova^ lung tumors 3 weeks post-CAR-T infusion. H) Representative flow plots and number of NK1.1^+^ NK cells in KP^ROR1-Ova^ lung tumors 3 weeks post-CAR-T infusion. I) CD69 expression on NK1.1^+^ NK cells in KP^ROR1-Ova^ lung tumors 3 weeks post-CAR-T infusion. J, K) Representative flow plots (J) and summary (K) of CD11b and CD27 expression on NK1.1^+^ NK cells in KP^ROR1-Ova^ lung tumors 3 weeks post-CAR-T infusion. All graphs: Mean +/- SD. One-way ANOVA with Tukey’s post-test. N=8-10 mice per group.

While CAR-T + NKTR-255 treatment enhanced DC activation in tumors, we also considered the possibility that a more inflammatory environment could simultaneously activate inhibitory pathways. In particular, PD-L1 is known to be upregulated in response to IFNγ,^79^ which we found was produced at higher levels by CAR-T cells after NKTR-255 treatment (Fig. 2F-H). Indeed, NKTR-255 + CAR-T combination therapy induced a significant increase in PD-L1 expression on both cDC1 and cDC2 in tumors to a greater extent than either therapy alone (Fig. 4E, 4F). Recent studies have demonstrated that co-expression of CD40, CD86, and PD-L1 is a feature of a subset of immunoregulatory DCs termed “mregDCs” (“mature dendritic cells enriched in maturation and regulatory molecules”), which co-localize with and maintain endogenous Tpex cells in mouse and human tumors.^80–82^ CAR-T + NKTR-255 treatment synergistically increased the number of CD40^+^CD86^+^PD-L1^+^ mregDC-like cells in lung tumors (Fig. 4G), correlating with the increased number of Endog. CD8^+^Tet^+^ Tpex cells observed in this group (Fig. 3L, 3M). Thus, CAR-T and NKTR-255 combination synergistically increases the number of activated DCs and mregDC-like cells in the tumor microenvironment, which correlates with increases in endogenous tumor-specific Tpex.

We next examined how CAR-T and/or NKTR-255 therapy impacted DCs in TdLNs, as migration and activation of tumor antigen-presenting DCs to TdLNs can also impact endogenous tumor-specific T cells.^83^ In addition to subsetting cDCs as cDC1 or cDC2, cDCs in lymph nodes can also be segregated as migratory or resident DCs based on their expression of MHC-II.^84^ While CAR-T monotherapy drove only a modest but insignificant increase in total cDC1 and cDC2 numbers within TdLNs (Fig. S3B), it significantly increased numbers of migratory cDC1s and cDC2s (Fig. S3B) and CD40^+^CD86^+^ “activated” cDC1s and cDC2 (Fig. S3D-F). This correlated with an increase in the number of Endog. CD8^+^Tet^+^ in the TdLN (Fig. S3G), suggestive of increased endogenous T cell priming. Surprisingly, addition of NKTR-255 to CAR-T therapy significantly decreased total numbers of cDC1 and cDC2, as well as migratory and CD40^+^CD86^+^ “activated” cDC1 and cDC2 in TdLNs compared to CAR-T therapy alone (Fig. S3B-F). These changes correlated with a lack of increase in Endog. CD8^+^Tet^+^ cells in the TdLN of CAR-T + NKTR-255 treated mice compared with CAR-T monotherapy (Fig. S3G). Neither NKTR-255 nor CAR-T therapy, alone or together, significantly impacted the differentiation of Endog. CD8^+^Tet^+^ cells in TdLNs, as the majority of cells adopted a Tcf1^+^Tim3^-^ Tpex phenotype, consistent with previous work (Fig. S3H).^50^ Altogether, these data demonstrate that CAR-T monotherapy promotes DC activation and migration to TdLNs, which correlates with an increase in endogenous tumor-specific T cells in TdLNs; however, CAR-T monotherapy does not significantly impact the phenotype of DCs within tumors, and increases terminal differentiation of endogenous tumor-specific T cells. In contrast, CAR-T and NKTR-255 combination therapy primarily activates DCs and endogenous tumor-specific T cells at the tumor site, not in the TdLN.

### Combination NKTR-255 and CAR-T treatment increases NK cell number and promotes their cytotoxic phenotype

In addition to expanding CD8^+^ T cells, IL-15 has a critical role in promoting NK cell expansion and development,^26^ and NKTR-255 has been shown to potently expand and differentiate NK cells at homeostasis and in models of hematologic malignancies.^33,34^ In contrast to these studies, NKTR-255 monotherapy only modestly and insignificantly increased numbers of NK cells in lung tumors (Fig. 4H). However, combination of CAR-T + NKTR-255 dramatically increased numbers of NK cells within tumors to a greater extent than either therapy alone (Fig. 4H). NKTR-255 also significantly increased expression of the activation marker CD69 on tumor-infiltrating NK cells, both as a monotherapy and in combination with CAR-T therapy (Fig. 4I). Interestingly, NKTR-255 monotherapy primarily promoted accumulation of CD27^+^ NK cells, which possess the greatest capacity to produce cytokines; by contrast, CAR-T + NKTR-255 combination therapy instead promoted accumulation of CD11b^+^ NK cells, which exhibit the greatest cytolytic function (Fig. 4J, 4K).^85,86^ Thus, CAR-T and NKTR-255 combination therapy synergistically activates and increases numbers of NK cells with enhanced cytolytic function within tumors.

### NKTR-255 synergizes with CAR-T cells to eliminate hROR1^+^ tumors and extend survival

Given that CAR-T and NKTR-255 combination therapy significantly enhanced CAR-T accumulation and function, and synergistically activated DCs, NK cells, and endogenous T cells, we next asked whether this combination treatment enabled superior control of tumors compared to either therapy alone. We first transplanted mice with KP^ROR1-Ova^ tumors and treated them with NKTR-255 or vehicle and/or CAR-T therapy as before (Fig. 5A). NKTR-255 or CAR-T monotherapy slightly slowed tumor progression compared to vehicle-treated mice, with CAR-T therapy slightly more effective than NKTR-255 monotherapy (Fig. 5B). In contrast, NKTR-255 and CAR-T combination therapy significantly improved tumor control compared to either CAR-T or NKTR-255 monotherapy and was the only treatment to induce tumor regression (Fig. 5B). Mice treated with CAR-T or NKTR-255 monotherapy progressed with ROR1^+^ and H-2K^b^/SIINFEKL^+^ tumor, highlighting their inability to eliminate the CAR target and a model neoantigen, respectively. Only CAR-T and NKTR-255 combination therapy was able to induce complete elimination of ROR1^+^ and H-2K^b^/SIINFEKL^+^ tumor, such that only ROR1^-^H-2K^b^/SIINFEKL^-^ tumor progressed after transient tumor regression (Fig. 5C, 5D). Thus, combining NKTR-255 and CAR-T therapy induces superior tumor control to either monotherapy and enables complete elimination of tumors expressing the CAR target and model neoantigens.

**Figure 5.**
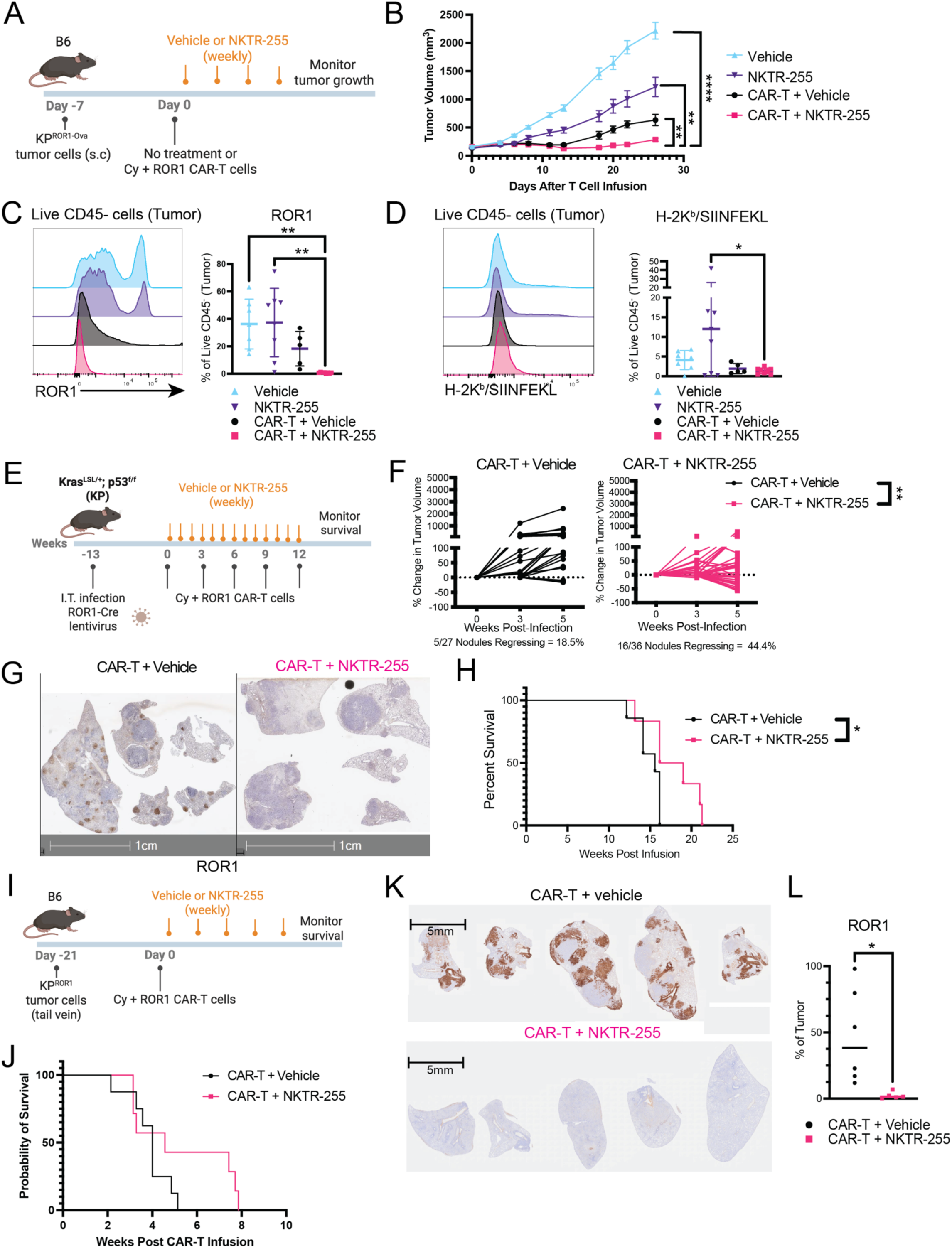
NKTR-255 synergizes with CAR-T cells to eliminate ROR1^+^ tumors and extend survival. A) Experimental schematic. Ova = ovalbumin. S.c. = subcutaneous. Cy = cyclophosphamide. B) Growth of KP^ROR1-Ova^ tumors in B6 mice treated as indicated. Mean +/- SEM. Two-way ANOVA with Holm-Šidák correction. N=7-8 mice per group. C, D) ROR1 (C) and H-2K^b^/SIINFEKL (D) expression on live CD45^-^ KP^ROR1-Ova^ tumor cells. Mean +/- SD. One-way ANOVA with Tukey’s post-test. N=7-8 mice per group. E) Experimental schematic. I.T. = intratracheal. Cy = cyclophosphamide. F) Percent change in KP^ROR1^ lung tumor volume measured by z-stack MRI. Lines indicate individual lung tumor nodules. Two-way ANOVA with Holm-Šidák correction. N=27-36 tumor nodules per group from N=6 mice per group G) Representative immunohistochemistry of ROR1 expression in KP^ROR1^ lung tumors at endpoint. H) Survival of KP^ROR1^ mice treated as indicated. Log-rank Mantel-Cox test. N=6-7 mice per group. I) Experimental schematic. Cy = cyclophosphamide. J) Survival of B6 mice transplanted with KP^ROR1^ lung tumors and treated as indicated. Log-rank Mantel-Cox test. N=7-8 mice per group. K) Quantification of ROR1^+^ cells in KP^ROR1^ lung tumors at endpoint. Mean shown. Two-way unpaired Student’s t-test. N=5-6 mice per group.

Ovalbumin is a strongly immunogenic model neoantigen that may induce inflammation in tumors that facilitates response to CAR-T and NKTR-255 therapy.^38^ Thus, to determine whether CAR-T and NKTR-255 combination therapy was similarly effective in “colder” tumors with lower mutational burden and neoantigen expression, we performed an identical experiment in the autochthonous KP^ROR1^ model, which we have previously shown develops “cold” tumors devoid of endogenous T cells, and monitored lung tumor growth by MRI (Fig 5E).^5,87^ NKTR-255 significantly slowed tumor progression over 5 weeks following a single infusion of CAR-T cells, inducing regression of ∼44.4% of tumor nodules compared to ∼18.5% of tumor nodules with CAR-T therapy alone (Fig. 5F). CAR-T and NKTR-255 therapy enabled complete elimination of ROR1^+^ tumor compared to CAR-T treatment alone, which resulted in significantly improved survival of KP^ROR1^ mice, with a subset of ∼50% of mice showing an extension in survival by ∼5 weeks (Fig. 5G, 5H). We observed similar effects in a model of ROR1^+^ lung metastasis, where a single infusion of CAR-T cells followed by weekly administration of NKTR-255 enabled complete elimination of ROR1^+^ tumor, and resulted in improved survival in a subset of ∼50% of mice (Fig. 5I-5L). Thus, combination of NKTR-255 with ROR1 CAR-T cells enables complete elimination of ROR1^+^ tumors, enhances tumor control, and improves survival in models of both “hot” and “cold” tumors. However, progression of ROR1^-^ tumor ultimately leads to tumor outgrowth after CAR-T and NKTR-255 treatment, limiting the magnitude of the survival benefit.

### Anti-tumor efficacy of NKTR-255 and CAR-T combination treatment depends on endogenous T cells

Given that CAR-T and NKTR-255 combination therapy significantly activated endogenous tumor-specific T cells in addition to CAR-T cells, it was unclear to what degree endogenous T cells contributed to the enhanced tumor control observed after combination therapy. To determine the role of endogenous T cells in controlling tumor growth, we implanted Thy1.2^+^ mice with KP^ROR1-Ova^ tumors, treated them with Cy + Thy1.1^+^ ROR1 CAR-T cells and administered either vehicle or NKTR-255 weekly thereafter (Fig. 6A). To deplete endogenous T cells, we used a previously described approach in which mice are treated with anti-Thy1.2 antibody to deplete endogenous Thy1.2^+^ T cells while sparing Thy1.1^+^ CAR-T cells.^43^ We confirmed that anti-Thy1.2 treatment eliminated endogenous T cells and did not drive a compensatory increase in CAR-T cells in the blood of treated mice (Fig. S4). As we observed before, NKTR-255 + CAR-T combination treatment significantly slowed tumor growth compared to CAR-T treatment alone (Fig. 6B, 6C). Depletion of Thy1.2 endogenous T cells slightly impaired tumor control by CAR-T cells, suggesting endogenous T cells play a small role in control of KP^ROR1-Ova^ tumors during CAR-T monotherapy. In contrast, Thy1.2 blockade significantly reduced tumor control by CAR-T + NKTR-255 combination therapy, to rates comparable to CAR-T treatment alone. To determine whether endogenous T cells play a functional role in tumor control in models that lack model neoantigens or high mutational burdens, we performed an identical experiment in a model of ROR1^+^ lung metastasis that lacked expression of Ova (Fig. 6D). In this model, Thy1.2 depletion had no impact on survival of mice treated with CAR-T monotherapy but dramatically reduced survival in the context of CAR-T + NKTR-255 combination therapy (Fig. 6E). Altogether, these data demonstrate that endogenous T cells play a critical role in mediating improved tumor control following CAR-T and NKTR-255 combination therapy.

**Figure 6.**
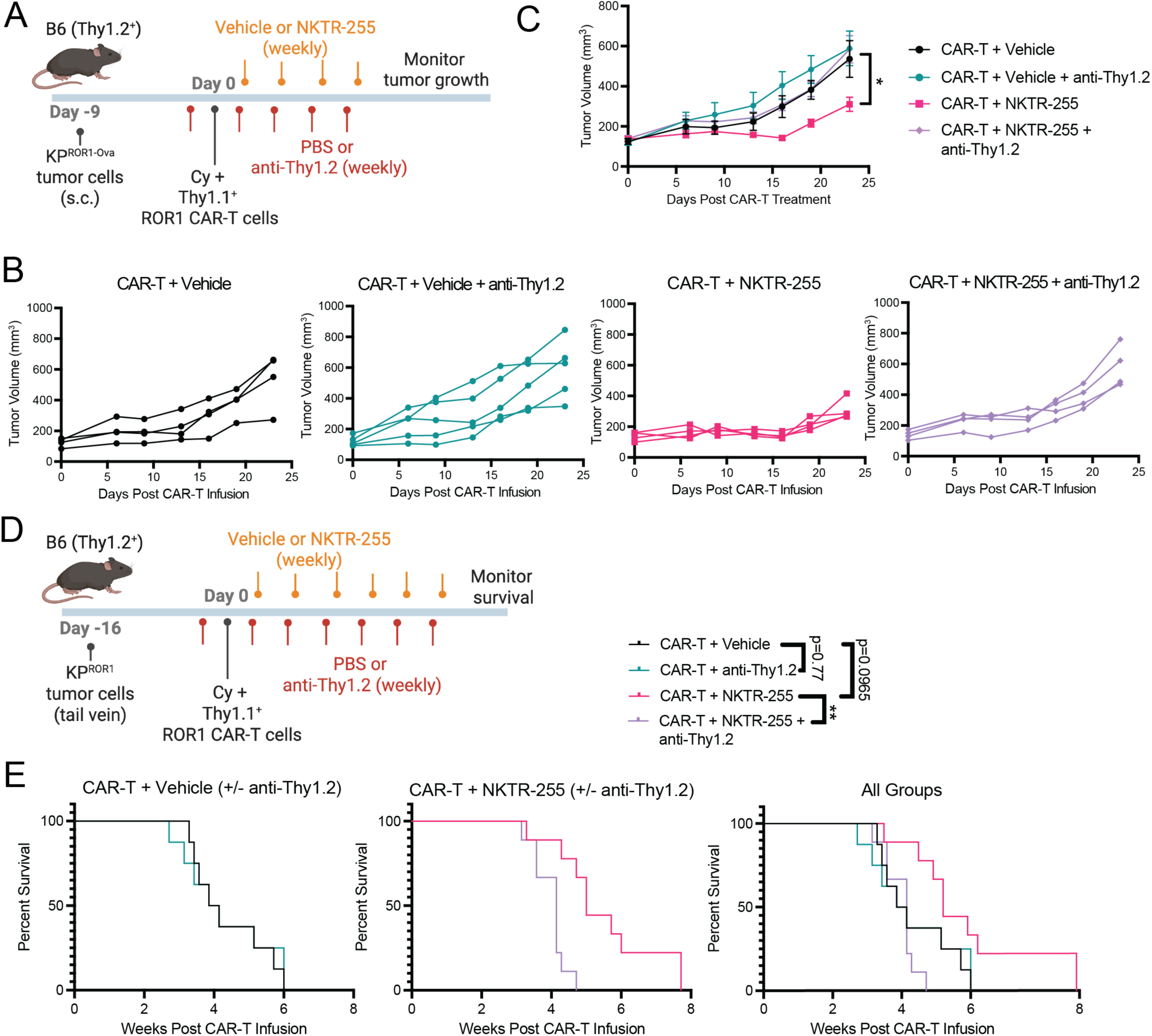
Anti-tumor efficacy of NKTR-255 and CAR-T combination treatment depends on endogenous T cells. A) Experimental schematic. Ova = ovalbumin. s.c. = subcutaneous. Cy = cyclophosphamide. B) KP^ROR1-Ova^ tumor volume in B6 mice treated as indicated. C) KP^ROR1-Ova^ tumor volume in (B) summarized across treatment groups. Mean +/- SEM. Two-way ANOVA with Holm-Šidák correction. N=4-5 mice per group. D) Experimental schematic. Cy = cyclophosphamide. E) Survival of B6 mice transplanted with KP^ROR1^ lung tumors and treated as indicated. Log-rank Mantel-Cox test. N=10 mice per group.

## Discussion

Here, we show that NKTR-255 significantly boosts accumulation, reduces exhaustion, and improves function of tumor-infiltrating CAR-T cells, enabling complete eradication of ROR1^+^ tumor in aggressive models of primary and metastatic lung cancer. In addition to increasing CD11b^+^ cytotoxic NK cells and activated CD40^+^CD86^+^ DCs within tumors, combination therapy with CAR-T cells and NKTR-255 also induced a synergistic increase in endogenous tumor-specific T cells that preserved a PD-1^+^Tcf1^+^ stem-like phenotype within tumors. These effects on endogenous T cells played a critical role in therapeutic efficacy, as improved tumor control after CAR-T and NKTR-255 treatment was almost completely ablated in the absence of endogenous T cells. Thus, our results demonstrate that combination of CAR-T and NKTR-255 therapy can favorably remodel the TME to promote both CAR-T-mediated and endogenous anti-tumor immunity greater than either monotherapy alone.

Though NKTR-255 has been shown to expand numbers of CD19 CAR-T cells in both preclinical models and patients with B cell malignancies,^29,35–37^ how it impacts the development of CAR-T cell dysfunction within solid tumors has not been previously examined. Our data show that NKTR-255 expands ROR1 CAR-T cell numbers, both systemically in the blood/spleen and within lung tumors, while preserving a less differentiated and more polyfunctional phenotype. This may be driven in part by selective expansion of less differentiated subsets, as our data show that PD-1^-^ CAR-T cells increased in number more dramatically than PD-1^+^ CAR-T cells upon NKTR-255 treatment. Indeed, PD-1^-^ T cells have been shown to express higher levels of the common-gamma chain of the IL-15 receptor (CD132) and be more sensitive to IL-15 *ex vivo*, suggesting less differentiated PD-1^-^ CAR-T cells may likewise preferentially expand in response to NKTR-255 treatment.^30^ Our data also suggest that NKTR-255 alters the phenotype of antigen-experienced PD-1^+^ CAR-T cells within tumors, suggesting NKTR-255 may protect CAR-T cells from terminal exhaustion. PD-1^+^ CAR-T cells in tumors of NKTR-255-treated mice showed reduced co-expression of LAG-3 and Tox.^59–62^ Recent studies have demonstrated that STAT5, the major mediator of IL-2 and IL-15 signaling, antagonizes Tox-dependent epigenetic programming of the exhausted state and instead fosters effector-like differentiation.^88,89^ Consistent with this, CAR-T cells from NKTR-255 treated mice showed significantly enhanced capacity to co-produce multiple effector cytokines and upregulated IFNγ response and cytotoxicity gene signatures. NKTR-255 also significantly reduced upregulation of Foxp3 and multiple genes associated with Foxp3^+^ Tregs. Foxp3 upregulation in CD8^+^ and CD4^+^ CAR-T cells may be driven by chronic CAR/TCR stimulation and/or TGFβ, which is highly expressed in human and mouse models of NSCLC.^68,90–92^ Indeed, CAR-T cells have been observed to upregulate Foxp3 in patients and show reduced cytotoxicity, increased suppressive activity, and correlate with disease progression.^66^ Thus, NKTR-255 not only boosts CAR-T cell numbers but protects their functional capacity in the suppressive lung TME.

Our data also demonstrate that endogenous T cells contribute significantly to tumor control in the context of CAR-T and NKTR-255 therapy, as improved efficacy was almost completely eliminated in the absence of endogenous T cells. NKTR-255 and CAR-T cell combination treatment synergistically activated endogenous tumor-specific Tpex cells within tumors, which promoted superior tumor control compared to either monotherapy. These activating effects on endogenous T cells were surprising, as they occurred despite the administration of lymphodepleting chemotherapy with Cy prior to CAR-T infusion. While Cy transiently ablates endogenous T cells, several studies suggest it can have immune stimulating effects that promote anti-tumor immunity, including through selective depletion of Tregs,^93–95^ enhanced activation and recruitment of tumor antigen-presenting DCs,^96,97^ and upregulation of pro-inflammatory cytokines and chemokines.^98,99^ Thus, administration of Cy with NKTR-255 could facilitate the recovery of endogenous tumor-specific Tpex cells, which are associated with sustained anti-tumor immunity.^75,100,101^ Indeed, emerging clinical evidence suggests that endogenous T cell activation and differentiation state correlates with durable CAR-T cell responses in patients. In a clinical trial evaluating CD19 CAR-T cells and anti-PD-L1 in patients with large B cell lymphoma (**LBCL**), on-going anti-PD-L1 treatment was associated with late and durable regression of both CD19^+^ and CD19^-^ tumor cells, suggesting a role for anti-PD-L1 in promoting endogenous T cell activity against CD19-negative tumor cells.^102^ Long-term responders to BCMA CAR-T therapy also show endogenous T cells with greater TCR diversity and less differentiated phenotypes, suggesting endogenous stem-like T cells may contribute to durable tumor control.^103^ Enhancing IL-15 signaling may enhance these axes, as a recent study demonstrated that IL-15-secreting CAR-T cells targeting GPC3 induced improved clinical responses that correlated with greater clonal expansion amongst CAR^-^ tumor-infiltrating T cells.^21^ Additionally, preliminary results from a phase 1b trial at our Center evaluating NKTR-255 in combination with CD19 CAR-T therapy demonstrated improved responses in LBCL patients that correlated with re-expansion of both CAR-T cells and stem-like CAR-negative T cells.^104^ While it is unclear whether the CAR-negative T cells expanded in these patients are tumor-reactive and contribute to tumor control, our data in mouse models provide functional evidence that CAR-T and NKTR-255 therapy together effectively expand endogenous tumor-specific T cells that play a major role in mediating therapeutic efficacy.

Although NKTR-255 significantly enhanced both CAR-T and endogenous T cell anti-tumor activity, all mice eventually succumbed to progression of ROR1^-^ tumor, indicating that additional strategies are needed to prevent tumor escape. CAR-T cell-mediated activation of endogenous T cells specific for distinct tumor antigens has been shown to enhance control of tumors with heterogeneous CAR target expression and prevent tumor escape in preclinical models.^18,19,76^ While our data show that endogenous T cells contributed significantly to tumor control after CAR-T and NKTR-255 treatment, tumors that lacked expression of both ROR1 and the model neoantigen Ova still progressed, indicating that endogenous T cell activation was insufficient to control tumor escape. Incomplete control of ROR1^-^ tumors could be due to the low mutational burden of KP tumors, which may fail to generate additional neoantigens targetable by endogenous T cells.^87^ Another possible mechanism driving tumor escape suggested by our data is insufficient engagement of TdLNs by CAR-T and NKTR-255 therapy. Epitope spreading canonically involves activation and migration of DCs presenting tumor antigens to TdLNs, where they can prime endogenous T cells specific for distinct tumor antigens.^105^ Whereas CAR-T cell monotherapy increased migratory DCs and tumor-specific T cells in TdLN, combination of CAR-T therapy with NKTR-255 preferentially increased endogenous T cell and DC activation within tumors but decreased their accumulation within TdLNs. These data suggest that CAR-T and NKTR-255 combination therapy may locally reactivate tumor-specific T cells within tumors rather than induce priming of new tumor-specific T cells in TdLNs. Strategies to promote DC maturation and migration to TdLNs, for example via co-administration of TLR or STING agonists,^106,107^ could potentially broaden CAR-T and NKTR-255-induced epitope spreading and combat escape of ROR1^-^ tumor more effectively.

Altogether, our data suggest that combining NKTR-255 with CAR-T therapy is a promising strategy to enhance both CAR-T and endogenous anti-tumor immunity to promote coordinated control of aggressive tumors, overcoming many of the major obstacles to CAR-T cell efficacy in solid tumors.

## Acknowledgements

We wish to thank the Benaroya Research Institute Genomics Core (RRID:SCR_026658) for bulk RNA-sequencing. The M.J. Murdock Charitable Trust generously supported the equipment used by this core facility. This research was also supported by the Flow Cytometry, Preclinical Imaging, Comparative Medicine, Experimental Histopathology, and the Genomics & Bioinformatics Shared Resources of the Fred Hutch/University of Washington/Seattle Children’s Cancer Consortium (P30 CA015704). Treatment schematic figures were created with BioRender.

## Funding

National Cancer Institute, R37CA276285 (S.S.)

American Lung Association, Lung Cancer Discovery Award (S.S.)

Nektar Therapeutics, Sponsored Research Agreement (S.S.)

American Cancer Society, Research Scholar Grant (S.S.)

V Foundation, Bob Bast Translational Research Grant (S.S.)

Cancer Center Support Grant, P30 CA015704 (S.S.)

National Institutes of Health, F31CA294918 (W.S.N.)

University of Washington ITHS, TL1 TR002318 (W.S.N.)

National Institutes of Health, F30CA305905 (M.G.K.)

## Author Contributions

Conceptualization: WSN, SS

Methodology: WSN, KB, MH, MM, SS

Investigation: WSN, MGK, EB, AJS, EG, DM, VZ, SM, SL, MH, LL, MV, SG, MS

Visualization: WSN, SS

Funding acquisition: WSN, MGK, SS

Project administration: WSN, SS

Supervision: SS

Writing – original draft: WSN, SS

Writing – review & editing: WSN, MGK, AJS, SM, MM, SS

## References

1 Patel, K. K., Tariveranmoshabad, M., Kadu, S., Shobaki, N. & June, C. From concept to cure: The evolution of CAR-T cell therapy. Molecular Therapy (2025). 10.1016/j.ymthe.2025.03.005

2 Du, B. et al. CAR-T therapy in solid tumors. Cancer Cell 43, 665–679 (2025). 10.1016/j.ccell.2025.03.019

3 Albelda, S. M. CAR T cell therapy for patients with solid tumours: key lessons to learn and unlearn. Nature Reviews Clinical Oncology 21, 47–66 (2024). 10.1038/s41571-023-00832-4

4 Jaeger-Ruckstuhl, C. A. et al. Phase 1 Study of ROR1 Specific CAR T Cells in Advanced Hematopoietic and Epithelial Malignancies. Clin Cancer Res (2024). 10.1158/1078-0432.CCR-24-2172

5 Srivastava, S. et al. Immunogenic Chemotherapy Enhances Recruitment of CAR-T Cells to Lung Tumors and Improves Antitumor Efficacy when Combined with Checkpoint Blockade. Cancer Cell 39, 193–208 e110 (2021). 10.1016/j.ccell.2020.11.005

6 Majzner, R. G. & Mackall, C. L. Tumor Antigen Escape from CAR T-cell Therapy. Cancer discovery (2018). 10.1158/2159-8290.cd-18-0442

7 Lin, H., Yang, X., Ye, S., Huang, L. & Mu, W. Antigen escape in CAR-T cell therapy: Mechanisms and overcoming strategies. Biomed Pharmacother 178, 117252 (2024). 10.1016/j.biopha.2024.117252

8 Schultz, L. M. et al. Outcomes After Nonresponse and Relapse Post-Tisagenlecleucel in Children, Adolescents, and Young Adults With B-Cell Acute Lymphoblastic Leukemia. J Clin Oncol 41, 354–363 (2023). 10.1200/jco.22.01076

9 Plaks, V. et al. CD19 target evasion as a mechanism of relapse in large B-cell lymphoma treated with axicabtagene ciloleucel. Blood 138, 1081–1085 (2021). 10.1182/blood.2021010930

10 Neelapu, S. S. et al. Axicabtagene ciloleucel CAR T-cell therapy in refractory large B-cell lymphoma. New England Journal of Medicine 377, 2531–2544 (2017).

11 Shah, B. D. et al. KTE-X19 for relapsed or refractory adult B-cell acute lymphoblastic leukaemia: phase 2 results of the single-arm, open-label, multicentre ZUMA-3 study. Lancet 398, 491–502 (2021). 10.1016/s0140-6736(21)01222-8

12 O’Rourke, D. M. et al. A single dose of peripherally infused EGFRvIII-directed CAR T cells mediates antigen loss and induces adaptive resistance in patients with recurrent glioblastoma. Science translational medicine 9, eaaa0984 (2017). 10.1126/scitranslmed.aaa0984

13 Zhong, G. et al. Complete remission of advanced pancreatic cancer induced by claudin18.2-targeted CAR-T cell therapy: a case report. Front Immunol 15, 1325860 (2024). 10.3389/fimmu.2024.1325860

14 Gargett, T. et al. Safety and biological outcomes following a phase 1 trial of GD2-specific CAR-T cells in patients with GD2-positive metastatic melanoma and other solid cancers. J Immunother Cancer 12 (2024). 10.1136/jitc-2023-008659

15 Beatty, G. L. et al. Activity of Mesothelin-Specific Chimeric Antigen Receptor T Cells Against Pancreatic Carcinoma Metastases in a Phase 1 Trial. Gastroenterology 155, 29–32 (2018). 10.1053/j.gastro.2018.03.029

16 Brossart, P. The Role of Antigen Spreading in the Efficacy of Immunotherapies. Clin Cancer Res 26, 4442–4447 (2020). 10.1158/1078-0432.Ccr-20-0305

17 Lu, Y. & Zhao, F. Strategies to overcome tumour relapse caused by antigen escape after CAR T therapy. Molecular Cancer 24, 126 (2025). 10.1186/s12943-025-02334-6

18 Conde, E. et al. Epitope spreading driven by the joint action of CART cells and pharmacological STING stimulation counteracts tumor escape via antigen-loss variants. J Immunother Cancer 9 (2021). 10.1136/jitc-2021-003351

19 Ma, L. et al. Vaccine-boosted CAR T crosstalk with host immunity to reject tumors with antigen heterogeneity. Cell 186, 3148–3165.e3120 (2023). 10.1016/j.cell.2023.06.002

20 Adusumilli, P. S. et al. A Phase I Trial of Regional Mesothelin-Targeted CAR T-cell Therapy in Patients with Malignant Pleural Disease, in Combination with the Anti-PD-1 Agent Pembrolizumab. Cancer Discov 11, 2748–2763 (2021). 10.1158/2159-8290.Cd-21-0407

21 Steffin, D. et al. Interleukin-15-armoured GPC3 CAR T cells for patients with solid cancers. Nature 637, 940–946 (2025). 10.1038/s41586-024-08261-8

22 Wang, L. et al. Expansion of endogenous T cells in CSF of pediatric CNS tumor patients undergoing locoregional delivery of IL13R〿2-targeting CAR T cells: an interim analysis. Res Sq (2023). 10.21203/rs.3.rs-3454977/v1

23 Chapuis, A. G. et al. T-Cell Therapy Using Interleukin-21-Primed Cytotoxic T-Cell Lymphocytes Combined With Cytotoxic T-Cell Lymphocyte Antigen-4 Blockade Results in Long-Term Cell Persistence and Durable Tumor Regression. J Clin Oncol (2016). 10.1200/JCO.2015.65.5142

24 Marks-Konczalik, J. et al. IL-2-induced activation-induced cell death is inhibited in IL-15 transgenic mice. Proc Natl Acad Sci U S A 97, 11445–11450 (2000). 10.1073/pnas.200363097

25 Judge, A. D., Zhang, X., Fujii, H., Surh, C. D. & Sprent, J. Interleukin 15 controls both proliferation and survival of a subset of memory-phenotype CD8(+) T cells. J Exp Med 196, 935–946 (2002). 10.1084/jem.20020772

26 Zhou, Y. et al. Interleukin 15 in Cell-Based Cancer Immunotherapy. Int J Mol Sci 23 (2022). 10.3390/ijms23137311

27 Becker, T. C. et al. Interleukin 15 is required for proliferative renewal of virus-specific memory CD8 T cells. J Exp Med 195, 1541–1548 (2002). 10.1084/jem.20020369

28 Steel, J. C., Waldmann, T. A. & Morris, J. C. Interleukin-15 biology and its therapeutic implications in cancer. Trends Pharmacol Sci 33, 35–41 (2012). 10.1016/j.tips.2011.09.004

29 Hirayama, A. V. et al. A novel polymer-conjugated human IL-15 improves efficacy of CD19-targeted CAR T-cell immunotherapy. Blood Adv 7, 2479–2493 (2023). 10.1182/bloodadvances.2022008697

30 Lee, J. et al. IL-15 promotes self-renewal of progenitor exhausted CD8 T cells during persistent antigenic stimulation. Front Immunol 14, 1117092 (2023). 10.3389/fimmu.2023.1117092

31 Di Pilato, M. et al. CXCR6 positions cytotoxic T cells to receive critical survival signals in the tumor microenvironment. Cell 184, 4512–4530 e4522 (2021). 10.1016/j.cell.2021.07.015

32 Waldmann, T. A. The biology of interleukin-2 and interleukin-15: implications for cancer therapy and vaccine design. Nature Reviews Immunology 6, 595–601 (2006). 10.1038/nri1901

33 Miyazaki, T. et al. NKTR-255, a novel polymer-conjugated rhIL-15 with potent antitumor efficacy. J Immunother Cancer 9 (2021). 10.1136/jitc-2020-002024

34 Robinson, T. O. et al. NKTR-255 is a polymer-conjugated IL-15 with unique mechanisms of action on T and natural killer cells. J Clin Invest 131 (2021). 10.1172/jci144365

35 Ahmed, S. et al. NKTR-255 Vs Placebo to Enhance Complete Responses and Durability Following CD19-Directed CAR-T Therapy in Patients with Relapsed/ Refractory (R/R) Large B-Cell Lymphoma (LBCL). Blood 144, 2068–2068 (2024). 10.1182/blood-2024-203576

36 Srinagesh, H. et al. A phase 1 clinical trial of NKTR-255 with CD19-22 CAR T-cell therapy for refractory B-cell acute lymphoblastic leukemia. Blood 144, 1689–1698 (2024). 10.1182/blood.2024024952

37 Vinaud Hirayama, A., Chou, C., Maloney, D. G., Marcondes, M. Q. & Turtle, C. J. A Phase Ib Open-Label Study Evaluating the Safety and Efficacy of NKTR-255 in Combination with CD19-Directed CAR-T Cell Therapy in Patients with Relapsed/Refractory (R/R) Large B-Cell Lymphoma (LBCL). Blood 140, 12747–12748 (2022). 10.1182/blood-2022-160191

38 DuPage, M. et al. Endogenous T cell responses to antigens expressed in lung adenocarcinomas delay malignant tumor progression. Cancer cell 19, 72–85 (2011). 10.1016/j.ccr.2010.11.011

39 DuPage, M. & Jacks, T. Genetically engineered mouse models of cancer reveal new insights about the antitumor immune response. Current opinion in immunology 25, 192–199 (2013). 10.1016/j.coi.2013.02.005

40 Hudecek, M. et al. The B-cell tumor-associated antigen ROR1 can be targeted with T cells modified to express a ROR1-specific chimeric antigen receptor. Blood 116, 4532–4541 (2010). 10.1182/blood-2010-05-283309

41 Hudecek, M. et al. The nonsignaling extracellular spacer domain of chimeric antigen receptors is decisive for in vivo antitumor activity. Cancer immunology research 3, 125–135 (2015). 10.1158/2326-6066.cir-14-0127

42 Snyder, A. J. et al. Modulating AP-1 enables CAR-T cells to establish an intratumoral PD-1+Tcf1+ stem-like reservoir and overcomes resistance to PD-1 axis blockade. bioRxiv, 2025.2004.2010.648245 (2025). 10.1101/2025.04.10.648245

43 Walsh, S. R. et al. Endogenous T cells prevent tumor immune escape following adoptive T cell therapy. J Clin Invest 129, 5400–5410 (2019). 10.1172/jci126199

44 Dolton, G. et al. More tricks with tetramers: a practical guide to staining T cells with peptide–MHC multimers. Immunology 146, 11–22 (2015). 10.1111/imm.12499

45 DuPage, M., Dooley, A. L. & Jacks, T. Conditional mouse lung cancer models using adenoviral or lentiviral delivery of Cre recombinase. Nature protocols 4, 1064–1072 (2009). 10.1038/nprot.2009.95

46 Rhode, P. R. et al. Comparison of the Superagonist Complex, ALT-803, to IL15 as Cancer Immunotherapeutics in Animal Models. Cancer Immunol Res 4, 49–60 (2016). 10.1158/2326-6066.Cir-15-0093-t

47 Han, K. P. et al. IL-15:IL-15 receptor alpha superagonist complex: high-level co-expression in recombinant mammalian cells, purification and characterization. Cytokine 56, 804–810 (2011). 10.1016/j.cyto.2011.09.028

48 Anderson, K. G. et al. Intravascular staining for discrimination of vascular and tissue leukocytes. Nat Protoc 9, 209–222 (2014). 10.1038/nprot.2014.005

49 Pereira, J. P., An, J., Xu, Y., Huang, Y. & Cyster, J. G. Cannabinoid receptor 2 mediates the retention of immature B cells in bone marrow sinusoids. Nat Immunol 10, 403–411 (2009). 10.1038/ni.1710

50 Connolly, K. A. et al. A reservoir of stem-like CD8(+) T cells in the tumor-draining lymph node preserves the ongoing antitumor immune response. Sci Immunol 6, eabg7836 (2021). 10.1126/sciimmunol.abg7836

51 Alizadeh, D. et al. IL15 Enhances CAR-T Cell Antitumor Activity by Reducing mTORC1 Activity and Preserving Their Stem Cell Memory Phenotype. Cancer Immunol Res 7, 759–772 (2019). 10.1158/2326-6066.Cir-18-0466

52 Liu, R. et al. PD-1 signaling negatively regulates the common cytokine receptor gamma chain via MARCH5-mediated ubiquitination and degradation to suppress anti-tumor immunity. Cell Res 33, 923–939 (2023). 10.1038/s41422-023-00890-4

53 Blackburn, S. D., Shin, H., Freeman, G. J. & Wherry, E. J. Selective expansion of a subset of exhausted CD8 T cells by αPD-L1 blockade. Proc National Acad Sci 105, 15016–15021 (2008). doi:10.1073/pnas.0801497105

54 Kansy, B. A. et al. PD-1 Status in CD8(+) T Cells Associates with Survival and Anti-PD-1 Therapeutic Outcomes in Head and Neck Cancer. Cancer Res 77, 6353–6364 (2017). 10.1158/0008-5472.Can-16-3167

55 Ma, J. et al. PD1Hi CD8+ T cells correlate with exhausted signature and poor clinical outcome in hepatocellular carcinoma. Journal for ImmunoTherapy of Cancer 7, 331 (2019). 10.1186/s40425-019-0814-7

56 Thommen, D. S. et al. Progression of Lung Cancer Is Associated with Increased Dysfunction of T Cells Defined by Coexpression of Multiple Inhibitory Receptors. Cancer Immunology Research 3, 1344–1355 (2015). 10.1158/2326-6066.Cir-15-0097

57 Andrews, L. P. et al. LAG-3 and PD-1 synergize on CD8+ T cells to drive T cell exhaustion and hinder autocrine IFNg-dependent anti-tumor immunity. Cell 187, 4355–4372.e4322 (2024). 10.1016/j.cell.2024.07.016

58 Cillo, A. R. et al. Blockade of LAG-3 and PD-1 leads to co-expression of cytotoxic and exhaustion gene modules in CD8+ T cells to promote antitumor immunity. Cell 187, 4373–4388.e4315 (2024). 10.1016/j.cell.2024.06.036

59 Alfei, F. et al. TOX reinforces the phenotype and longevity of exhausted T cells in chronic viral infection. Nature 571, 265–269 (2019). 10.1038/s41586-019-1326-9

60 Khan, O. et al. TOX transcriptionally and epigenetically programs CD8+ T cell exhaustion. Nature 571, 211–218 (2019). 10.1038/s41586-019-1325-x

61 Scott, A. C. et al. TOX is a critical regulator of tumour-specific T cell differentiation. Nature 571, 270–274 (2019). 10.1038/s41586-019-1324-y

62 Seo, H. et al. TOX and TOX2 transcription factors cooperate with NR4A transcription factors to impose CD8+ T cell exhaustion. Proceedings of the National Academy of Sciences of the United States of America 116, 12410–12415 (2019). 10.1073/pnas.1905675116

63 Ciuffreda, D. et al. Polyfunctional HCV-specific T-cell responses are associated with effective control of HCV replication. Eur J Immunol 38, 2665–2677 (2008). 10.1002/eji.200838336

64 Almeida, J. R. et al. Superior control of HIV-1 replication by CD8+ T cells is reflected by their avidity, polyfunctionality, and clonal turnover. J Exp Med 204, 2473–2485 (2007). 10.1084/jem.20070784

65 Tang, N. et al. TGF-beta inhibition via CRISPR promotes the long-term efficacy of CAR T cells against solid tumors. JCI Insight 5 (2020). 10.1172/jci.insight.133977

66 Good, Z. et al. Post-infusion CAR T(Reg) cells identify patients resistant to CD19-CAR therapy. Nat Med 28, 1860–1871 (2022). 10.1038/s41591-022-01960-7

67 Peter, L. et al. Effects of CRISPR-Cas9-mediated FOXP3 knockout on CAR T cell potency. Molecular Therapy Methods & Clinical Development 33 (2025). 10.1016/j.omtm.2025.101570

68 Lozano, T. et al. TCR-induced FOXP3 expression by CD8(+) T cells impairs their anti-tumor activity. Cancer Lett 528, 45–58 (2022). 10.1016/j.canlet.2021.12.030

69 Kim, R. H. et al. Effect of chimeric antigen receptor (CAR) T cells on clonal expansion of endogenous non-CAR T cells in patients (pts) with advanced solid cancer. J Clin Oncol 35, 3011–3011 (2017). 10.1200/JCO.2017.35.15_suppl.3011

70 Hont, A. B. et al. 612 Lymphodepletion blunts the antigen-spreading response post TAA-T cell therapy for pediatric solid tumors. Journal for ImmunoTherapy of Cancer 11 (2023). 10.1136/jitc-2023-SITC2023.0612

71 Gros, A. et al. PD-1 identifies the patient-specific CD8(+) tumor-reactive repertoire infiltrating human tumors. J Clin Invest 124, 2246–2259 (2014). 10.1172/JCI73639

72 Fernandez-Poma, S. M. et al. Expansion of Tumor-Infiltrating CD8(+) T cells Expressing PD-1 Improves the Efficacy of Adoptive T-cell Therapy. Cancer Res 77, 3672–3684 (2017). 10.1158/0008-5472.Can-17-0236

73 Utzschneider, D. T. et al. Early precursor T cells establish and propagate T cell exhaustion in chronic infection. Nat Immunol 21, 1256–1266 (2020). 10.1038/s41590-020-0760-z

74 Kallies, A., Zehn, D. & Utzschneider, D. T. Precursor exhausted T cells: key to successful immunotherapy? Nat Rev Immunol 20, 128–136 (2020). 10.1038/s41577-019-0223-7

75 Miller, B. C. et al. Subsets of exhausted CD8(+) T cells differentially mediate tumor control and respond to checkpoint blockade. Nat Immunol 20, 326–336 (2019). 10.1038/s41590-019-0312-6

76 Lai, J. et al. Adoptive cellular therapy with T cells expressing the dendritic cell growth factor Flt3L drives epitope spreading and antitumor immunity. Nat Immunol 21, 914–926 (2020). 10.1038/s41590-020-0676-7

77 Mazzoccoli, L. & Liu, B. Dendritic Cells in Shaping Anti-Tumor T Cell Response. Cancers (Basel) 16 (2024). 10.3390/cancers16122211

78 Anguille, S. et al. Short-term cultured, interleukin-15 differentiated dendritic cells have potent immunostimulatory properties. J Transl Med 7, 109 (2009). 10.1186/1479-5876-7-109

79 Mandai, M. et al. Dual Faces of IFNγ in Cancer Progression: A Role of PD-L1 Induction in the Determination of Pro- and Antitumor Immunity. Clinical Cancer Research 22, 2329–2334 (2016). 10.1158/1078-0432.Ccr-16-0224

80 Maier, B. et al. A conserved dendritic-cell regulatory program limits antitumour immunity. Nature 580, 257–262 (2020). 10.1038/s41586-020-2134-y

81 Magen, A. et al. Intratumoral dendritic cell–CD4+ T helper cell niches enable CD8+ T cell differentiation following PD-1 blockade in hepatocellular carcinoma. Nature Medicine 29, 1389–1399 (2023). 10.1038/s41591-023-02345-0

82 Peng, Q. et al. PD-L1 on dendritic cells attenuates T cell activation and regulates response to immune checkpoint blockade. Nature Communications 11, 4835 (2020). 10.1038/s41467-020-18570-x

83 Pittet, M. J., Di Pilato, M., Garris, C. & Mempel, T. R. Dendritic cells as shepherds of T cell immunity in cancer. Immunity 56, 2218–2230 (2023). 10.1016/j.immuni.2023.08.014

84 Schenkel, J. M. et al. Conventional type I dendritic cells maintain a reservoir of proliferative tumor-antigen specific TCF-1(+) CD8(+) T cells in tumor-draining lymph nodes. Immunity 54, 2338–2353 e2336 (2021). 10.1016/j.immuni.2021.08.026

85 Fu, B. et al. CD11b and CD27 reflect distinct population and functional specialization in human natural killer cells. Immunology 133, 350–359 (2011). 10.1111/j.1365-2567.2011.03446.x

86 Zhang, Q.-F. et al. Liver-infiltrating CD11b−CD27− NK subsets account for NK-cell dysfunction in patients with hepatocellular carcinoma and are associated with tumor progression. Cellular & Molecular Immunology 14, 819–829 (2017). 10.1038/cmi.2016.28

87 McFadden, D. G. et al. Mutational landscape of EGFR-, MYC-, and Kras-driven genetically engineered mouse models of lung adenocarcinoma. Proc Natl Acad Sci U S A 113, E6409–e6417 (2016). 10.1073/pnas.1613601113

88 Ding, Z. C. et al. Persistent STAT5 activation reprograms the epigenetic landscape in CD4(+) T cells to drive polyfunctionality and antitumor immunity. Sci Immunol 5 (2020). 10.1126/sciimmunol.aba5962

89 Beltra, J. C. et al. Stat5 opposes the transcription factor Tox and rewires exhausted CD8(+) T cells toward durable effector-like states during chronic antigen exposure. Immunity 56, 2699–2718.e2611 (2023). 10.1016/j.immuni.2023.11.005

90 Chen, W. et al. Conversion of peripheral CD4+CD25-naive T cells to CD4+CD25+ regulatory T cells by TGF-beta induction of transcription factor Foxp3. J Exp Med 198, 1875–1886 (2003). 10.1084/jem.20030152

91 Li, J. et al. Prognostic value of TGF-beta in lung cancer: systematic review and meta-analysis. Bmc Cancer 19, 691 (2019). 10.1186/s12885-019-5917-5

92 Saito, A., Horie, M. & Nagase, T. TGF-beta Signaling in Lung Health and Disease. Int J Mol Sci 19 (2018). 10.3390/ijms19082460

93 Brode, S., Raine, T., Zaccone, P. & Cooke, A. Cyclophosphamide-induced type-1 diabetes in the NOD mouse is associated with a reduction of CD4+CD25+Foxp3+ regulatory T cells. J Immunol 177, 6603–6612 (2006). 10.4049/jimmunol.177.10.6603

94 Ghiringhelli, F. et al. CD4+CD25+ regulatory T cells suppress tumor immunity but are sensitive to cyclophosphamide which allows immunotherapy of established tumors to be curative. Eur J Immunol 34, 336–344 (2004). 10.1002/eji.200324181

95 Lutsiak, M. E. et al. Inhibition of CD4(+)25+ T regulatory cell function implicated in enhanced immune response by low-dose cyclophosphamide. Blood 105, 2862–2868 (2005). 10.1182/blood-2004-06-2410

96 Radojcic, V. et al. Cyclophosphamide resets dendritic cell homeostasis and enhances antitumor immunity through effects that extend beyond regulatory T cell elimination. Cancer Immunol Immunother 59, 137–148 (2010). 10.1007/s00262-009-0734-3

97 Nakahara, T. et al. Cyclophosphamide enhances immunity by modulating the balance of dendritic cell subsets in lymphoid organs. Blood 115, 4384–4392 (2010). 10.1182/blood-2009-11-251231

98 Murad, J. P. et al. Pre-conditioning modifies the TME to enhance solid tumor CAR T cell efficacy and endogenous protective immunity. Mol Ther 29, 2335–2349 (2021). 10.1016/j.ymthe.2021.02.024

99 Hughes, E. et al. T-cell modulation by cyclophosphamide for tumour therapy. Immunology 154, 62–68 (2018). 10.1111/imm.12913

100 Koh, J. et al. TCF1+PD-1+ tumour-infiltrating lymphocytes predict a favorable response and prolonged survival after immune checkpoint inhibitor therapy for non-small-cell lung cancer. European Journal of Cancer 174, 10–20 (2022). 10.1016/j.ejca.2022.07.004

101 Sade-Feldman, M. et al. Defining T Cell States Associated with Response to Checkpoint Immunotherapy in Melanoma. Cell 176, 404 (2019). 10.1016/j.cell.2018.12.034

102 Hirayama, A. V. et al. Timing of anti-PD-L1 antibody initiation affects efficacy/toxicity of CD19 CAR T-cell therapy for large B-cell lymphoma. Blood Adv 8, 453–467 (2024). 10.1182/bloodadvances.2023011287

103 Vieira dos Santos, J., et al. Long-term Remission After Cilta-Cel in Multiple Myeloma Is Linked to Diverse T Cells and Low Myeloid Suppression. Blood Advances (2025). 10.1182/bloodadvances.2025018078

104 Hirayama, A. V. et al. 139 | ENHANCED CAR T-CELL EXPANSION AND DURABLE COMPLETE RESPONSES WITH NKTR-255 PLUS LISOCABTAGENE MARALEUCEL IN RELAPSED/REFRACTORY LARGE B-CELL LYMPHOMA. Hematological Oncology 43, e139_70093 (2025). 10.1002/hon.70093_139

105 von Renesse, J., Lin, M.-C. & Ho, P.-C. Tumor-draining lymph nodes – friend or foe during immune checkpoint therapy? Trends in Cancer 11, 676–690 (2025). 10.1016/j.trecan.2025.04.008

106 Schmid, M. A., Takizawa, H., Baumjohann, D. R., Saito, Y. & Manz, M. G. Bone marrow dendritic cell progenitors sense pathogens via Toll-like receptors and subsequently migrate to inflamed lymph nodes. Blood 118, 4829–4840 (2011). 10.1182/blood-2011-03-344960

107 Wang, J. et al. STING licensing of type I dendritic cells potentiates antitumor immunity. Sci Immunol 9, eadj3945 (2024). 10.1126/sciimmunol.adj3945

