## Supplementary Figures and Tables for "NKTR-255, a polymer-conjugated IL-15, synergizes with CAR-T cell therapy to activate endogenous anti-tumor immunity and improve tumor control"

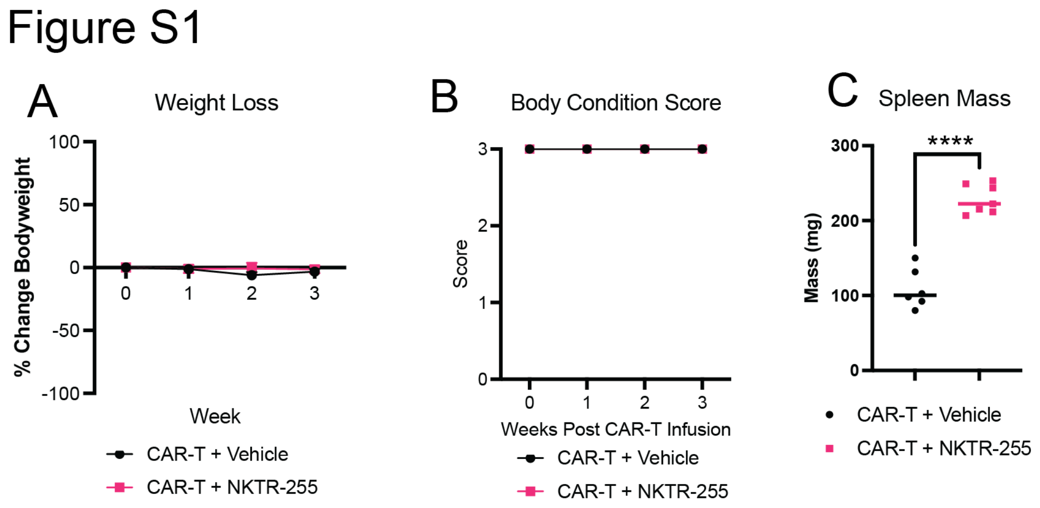


**Figure S1. CAR-T and NKTR-255 combination therapy is well-tolerated.** A, B) Percent weight loss (A) and body condition score (B) of KP$^{ROR1}$ mice treated as indicated. N=6-7 mice per group. C) Spleen mass of KP$^{ROR1}$ mice treated as indicated two weeks post-CAR-T cell infusion. Two-way unpaired Student's t-test. N=6-7 mice per group.

### Figure S2

A

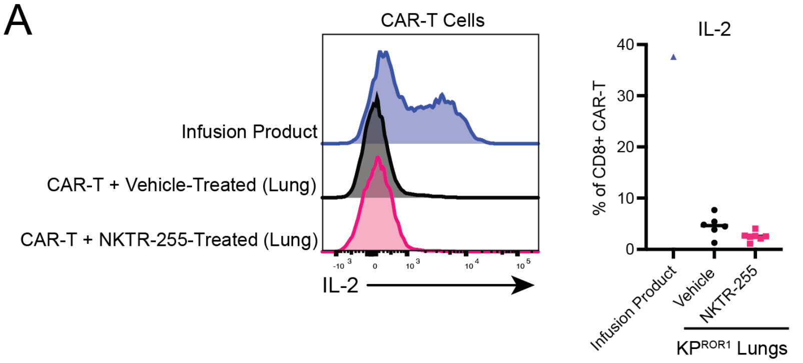


B

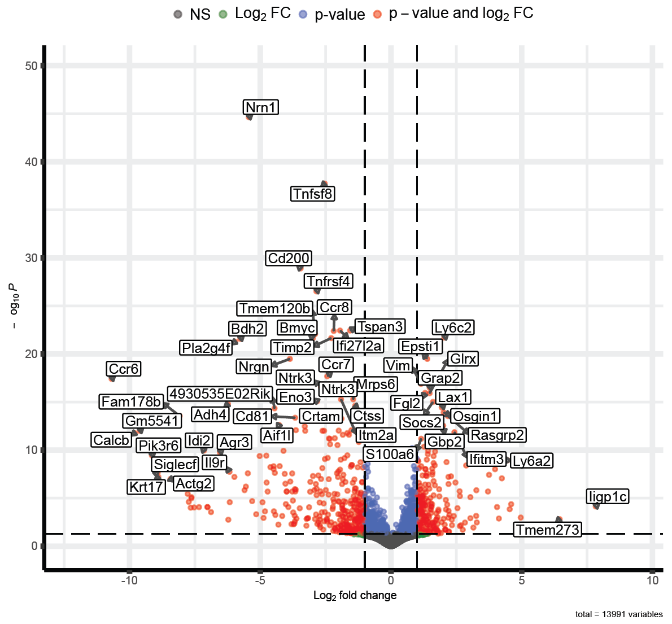


**Figure S2. Effects of NKTR-255 on ROR1 CAR-T cell phenotype.** A) IL-2 production by CD8$^+$CD19$^+$ CAR-T cells in KP$^{ROR1}$ lung tumors 2 weeks post-CAR-T cell infusion after *ex vivo* restimulation with PMA/ionomycin. N=6-7 mice per group. B) Volcano plot showing genes differentially expressed in CD8$^+$CD19$^+$ CAR-T cells sorted from lung tumors of KP$^{ROR1}$ mice treated with vehicle or NKTR-255 two weeks post-CAR-T cell infusion. N=6 mice per group

### Figure S3

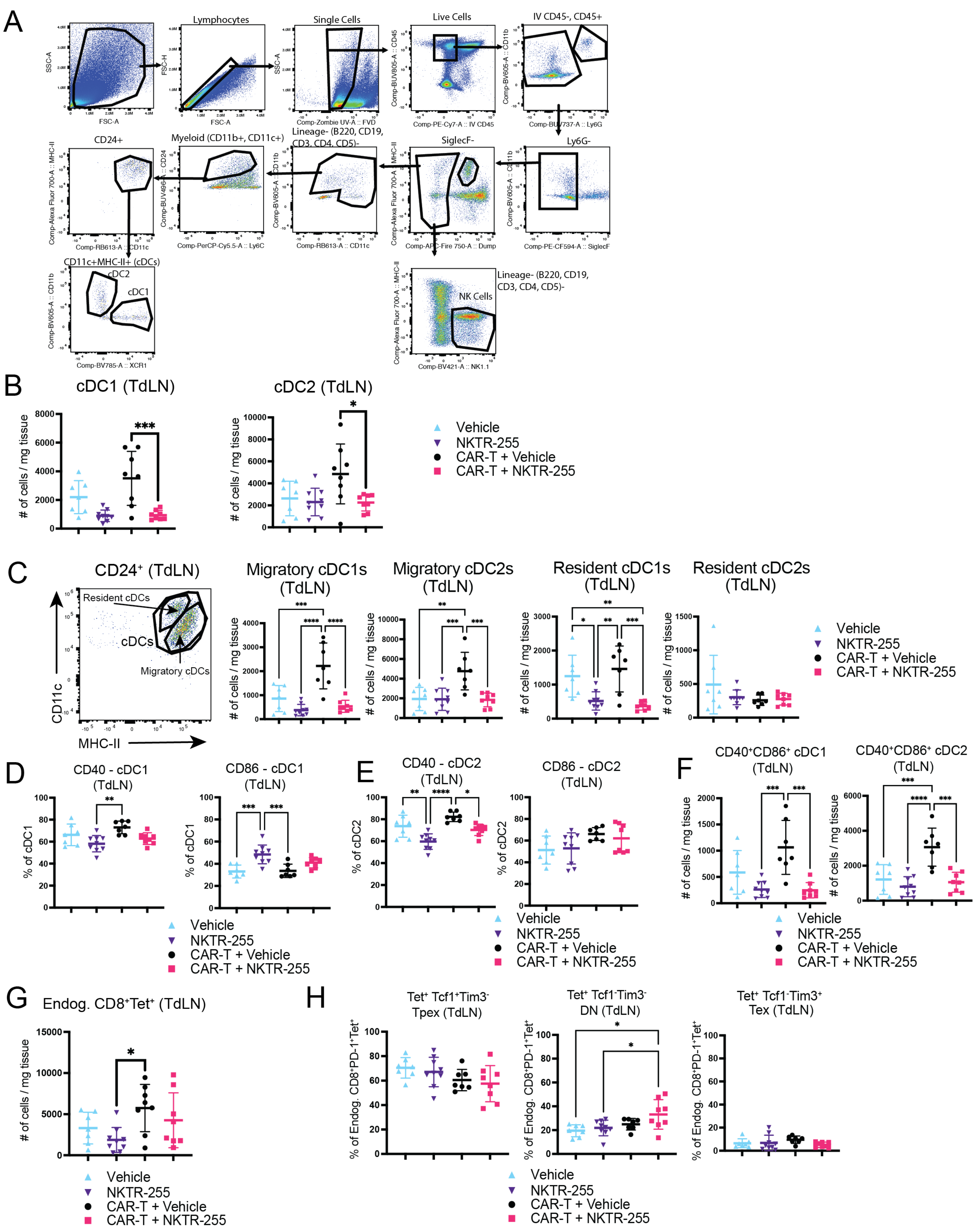
A
Lymphocytes
Single Cells
Live Cells
IV CD45-, CD45+
CD24+
Myeloid (CD11b+, CD11c+)
Lineage- (B220, CD19, CD3, CD4, CD5)-
SiglecF-
Ly6G-
CD11c+MHC-II+ (cDCs)
cDC2
cDC1
Lineage- (B220, CD19, CD3, CD4, CD5)-
NK Cells
B
cDC1 (TdLN)
cDC2 (TdLN)
### of cells / mg tissue
Vehicle
NKTR-255
CAR-T + Vehicle
CAR-T + NKTR-255
C
CD24⁺ (TdLN)
Resident cDCs
cDCs
Migratory cDCs
CD11c
MHC-II
Migratory cDC1s (TdLN)
Migratory cDC2s (TdLN)
Resident cDC1s (TdLN)
Resident cDC2s (TdLN)
D
CD40 - cDC1 (TdLN)
CD86 - cDC1 (TdLN)
% of cDC1
E
CD40 - cDC2 (TdLN)
CD86 - cDC2 (TdLN)
% of cDC2
F
CD40⁺CD86⁺ cDC1 (TdLN)
CD40⁺CD86⁺ cDC2 (TdLN)
G
Endog. CD8⁺Tet⁺ (TdLN)
H
Tet⁺ Tcf1⁺Tim3⁻ Tpex (TdLN)
Tet⁺ Tcf1⁻Tim3⁻ DN (TdLN)
Tet⁺ Tcf1⁻Tim3⁺ Tex (TdLN)
% of Endog. CD8⁺PD-1⁺Tet⁺

**Figure S3. NKTR-255 and ROR1 CAR-T cell therapy do not significantly impact DCs or endogenous T cells in tumor-draining lymph nodes.** A) Gating strategy for identification of cDC1, cDC2, and NK cells in lung tumors. B) Number of cDC1 and cDC2 in mediastinal tumor-draining lymph nodes (**TdLN**) of $KP^{ROR1\text{-}Ova}$ mice 3 weeks post-CAR-T cell infusion. C) Number of migratory and resident cDC1 and cDC2 in TdLN of $KP^{ROR1\text{-}Ova}$ mice 3 weeks post-CAR-T cell infusion. D, E) CD40 and CD86 expression on cDC1 (D) and cDC2 (E) in TdLN of $KP^{ROR1\text{-}Ova}$ mice 3 weeks post-CAR-T cell infusion. F) Number of $CD40^{+}CD86^{+}$ cDC1 and cDC2 in TdLN of $KP^{ROR1\text{-}Ova}$ mice 3 weeks post-CAR-T cell infusion. G) Number of endogenous $CD45.2^{+}$ H-$2K^{b}$/SIINFEKL tetramer$^{+}$ $CD8^{+}$ T cells in TdLN of $KP^{ROR1\text{-}Ova}$ mice 3 weeks post-CAR-T cell infusion. H) Frequency of $Tcf1^{+}Tim3^{-}$ Tpex, $Tcf1^{-}Tim3^{-}$ double negative (DN) and $Tcf1^{-}Tim3^{+}$ Tex among PD-$1^{+}$ endogenous $CD45.2^{+}$ H-$2K^{b}$/SIINFEKL tetramer$^{+}$ $CD8^{+}$ T cells in TdLN of $KP^{ROR1\text{-}Ova}$ mice 3 weeks post-CAR-T cell infusion. B-H: Mean +/- SD. One-way ANOVA with Tukey’s post-test. N=8-10 mice per group.

### Figure S4

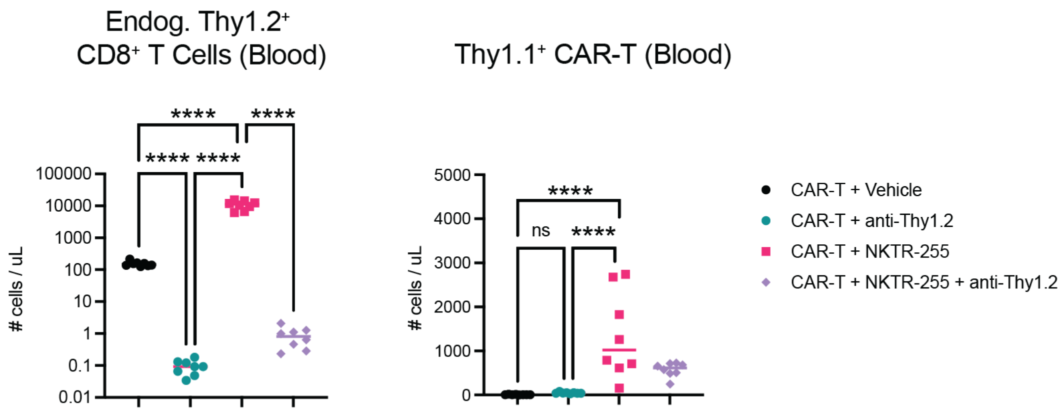


**Figure S4. Anti-Thy1.2 effectively depletes endogenous T cells in CAR-T and/or NKTR-255 treated mice.** Number of endogenous CD8$^{+}$Thy1.2$^{+}$ T cells (left) and CD8$^{+}$Thy1.1$^{+}$ CAR-T cells (right) in blood of mice bearing KP$^{ROR1\text{-}Ova}$ tumors 14 days post-CAR-T cell infusion. Mean shown. One-way ANOVA with Tukey's post-test. N=8 mice per group.

**Table S1. Flow antibodies used for immune profiling.**

**Flow panel for T cells in lungs, spleens, and TdLNs:**

| **Antibody Name** | **Dilution** | **Clone** | **Catalog Number** |
|---|---|---|---|
| CD8a BUV395 | 1:200 | 53-6.7 | BD 563786 |
| CD3e BUV496 | 1:100 | 145-2C11 | BD 612955 |
| CD45.2 BUV563 | 1:100 | 104 | BD 741273 |
| CD62L BUV737 | 1:100 | MEL-14 | BD 612833 |
| CD45.1 BUV805 | 1:100 | A20 | BD 741958 |
| CD44 BV510 | 1:100 | IM7 | BD 563114 |
| CD4 BV570 | 1:400 | RM-45 | BioLegend 100541 |
| Tim-3 BV605 | 1:200 | RMT3-23 | BioLegend 119721 |
| CX3CR1 BV785 | 1:100 | SA011F11 | BioLegend 149029 |
| CD19 FITC | 1:100 | eBio1D3 | Thermo 11-0193-82 |
| Lag-3 PE-Dazzle594 | 1:100 | C9157W | BioLegend 125223 |
| CD25 PE-Cy5 | 1:100 | PC61 | BioLegend 102010 |
| PD-1 APC eFluor750 | 1:100 | J43 | Thermo 47-9985-82 |
| Foxp3 eFluor450 | 1:100 | FJK-16s | Thermo 48-5773-82 |
| Ki-67 BV650 | 1:100 | B56 | BD 563757 |
| Tcf1 PE | 1:100 | C63D9 | Cell Signaling 14456 |
| Tox APC | 1:100 | REA473 | Miltenyi 130-118-335 |
| CD45 PE-Cy7 | 1:100 (infused i.v.) | 30-F11 | Biolegend 103114 |
| pMHC/SIIN Tetramer BV421 | 1:100 | | Made by Fred Hutch Core |
| Fc-ROR1 | 1:500 | | Acro RO1-H5250 |
| anti-Hu IgG Fc PerCP-Cy5.5 | 1:100 (secondary) | M1310405 | Biolegend 410710 |

**Flow panel for innate immune cells in lungs and TdLNs:**

| **Antibody Name** | **Dilution** | **Clone** | **Catalog Number** |
|---|---|---|---|
| CD103 BUV395 | 1:200 | 2E7 | BD 748253 |
| CD24 BUV496 | 1:100 | M1/69 | BD 612953 |
| CD27 BUV563 | 1:40 | LG.3A10 | BD 741275 |
| Ly6G BUV737 | 1:160 | 1A8 | BD 741813 |
| CD45 BUV805 | 1:320 | 30-F11 | BD 748370 |
| NK1.1 BV421 | 1:80 | PK136 | BioLegend 108731 |
| CD69 BV510 | 1:100 | H1.2F3 | BioLegend 104531 |
| CD11b BV605 | 1:100 | M1/70 | BioLegend 101237 |
| CD86 BV650 | 1:100 | GL-1 | BioLegend 105035 |
| CD8a BV711 | 1:100 | 53-6.7 | BioLegend 100747 |
| Xcr1 BV785 | 1:100 | ZET | BioLegend 148225 |

| | | | |
|---|---|---|---|
| F4/80 Alexa Fluor 488 | 1:100 | T45-2342 | BD 567201 |
| CD11c BB630 | 1:200 | N418 | BD Custom Conjugate |
| Ly6C PerCP-Cy5.5 | 1:200 | HK1.4 | BioLegend 128011 |
| H-2Kb SIINFEKL PE | 1:100 | 25-D1.16 | BioLegend 141604 |
| SiglecF PE-CF594 | 1:400 | E50-2440 | BD 562757 |
| CD40 PE-Cy5 | 1:100 | 3/23 | BioLegend 124617 |
| iNOS PE-Cy5.5 | 1:100 | CXNFT | Thermo eBio 35-5920-82 |
| PD-L1 APC | 1:100 | 10F.9G2 | BioLegend 124311 |
| I-A/I-E A700 | 1:400 | M5/114.15.2 | BioLegend 107622 |
| B220 APC/Fire 750 | 1:200 | RA3-6B2 | BioLegend 103259 |
| CD19 APC/Fire 750 | 1:200 | 6D5 | BioLegend 115557 |
| CD3e APC/Fire 750 | 1:200 | 145-2C11 | BioLegend 100361 |
| CD4 APC/Fire 750 | 1:200 | RM4-4 | BioLegend 116019 |
| CD5 APC/Fire 750 | 1:200 | 53-7.4 | BioLegend 100633 |
| CD45 PE-Cy7 | 1:100 (infused i.v.) | 30-F11 | Biolegend 103114 |

**Flow panels for blood:**

| **Antibody Name** | **Dilution** | **Clone** | **Catalog Number** |
|---|---|---|---|
| CD45.1 APC | 1:100 | A20 | BioLegend 110714 |
| CD45.2 BV711 | 1:100 | 104 | BioLegend 109847 |
| CD3 PE/Dazzle 594 | 1:100 | 17A2 | BioLegend 100246 |
| CD4 BV570 | 1:100 | RM4-5 | BioLegend 100541 |
| CD8 BV785 | 1:100 | 53-6.7 | BioLegend 100750 |
| CD19 PE-Cy7 | 1:100 | 6D5 | BioLegend 115520 |
| PD-1 APC eFluor780 | 1:100 | J43 | Thermo 47-9985-82 |
